# A constraint-based model reveals hysteresis in island biogeography

**DOI:** 10.1101/251926

**Authors:** Joseph R. Burger, Robert P. Anderson, Meghan A. Balk, Trevor S. Fristoe

## Abstract

**Aim:** We present a Constraint-based Model of Dynamic Island Biogeography (C-DIB) that predicts how species functional traits interact with dynamic environments to determine the candidate species available for local community assembly on real and habitat islands through time.

**Location:** Real and habitat islands globally.

**Methods:** We develop the C-DIB model concept, synthesize the relevant literature, and present a toolkit for evaluating model predictions for a wide variety of “island” systems and taxa.

**Results:** The C-DIB model reveals that as islands cycle between phases of increasing or decreasing size and connectivity to a source pool, the dominant process driving species’ presence or absence switches between colonization and extinction. Both processes are mediated by interactions between organismal traits and environmental constraints. Colonization probability is predicted by a species’ ability to cross the intervening matrix between a population source and the island; population persistence (or extinction) is predicted by the minimum spatial requirements to sustain an isolated population. The non-random distributions of mammals on islands of the Sunda Shelf and Great Basin “sky islands” provide example study systems for evaluating the C-DIB model.

**Main conclusions:** Because different suites of traits impose constraints on the processes of colonization and extinction, similar environmental conditions can host different candidate species despite the same predicted richness. Thus, the model exemplifies the specific yet underappreciated role of *hysteresis* –the dependency of outcomes not only on the current system state –but also the historical contingency of environmental change in affecting populations and communities in insular systems.

## INTRODUCTION

The earth is dynamic and many natural systems cycle in predictable ways. Landscapes change over space and time resulting in patches of habitats that expand and contract, appear and disappear, connect and disconnect. For example, cycles of variation in the Earth’s orbit, identified by Milankovich, drive global climate change over geological time scales. Throughout the Pleistocene, these cycles caused repeated sea-level changes as well as latitudinal and elevational shifts in climate, creating archipelagos of real and habitat islands that changed through time (Box 1a and b). Over shorter time periods at smaller geographic extents, natural and anthropogenic forces shape similar cycles as habitat patches are alternately fragmented and united by disturbance and succession (Box 1c). As fundamental features of the planet, these directional, cyclical, and predictable changes have undoubtedly played a strong role in affecting population and community dynamics, species distributions, and the generation and maintenance of biodiversity.

### Box 1.

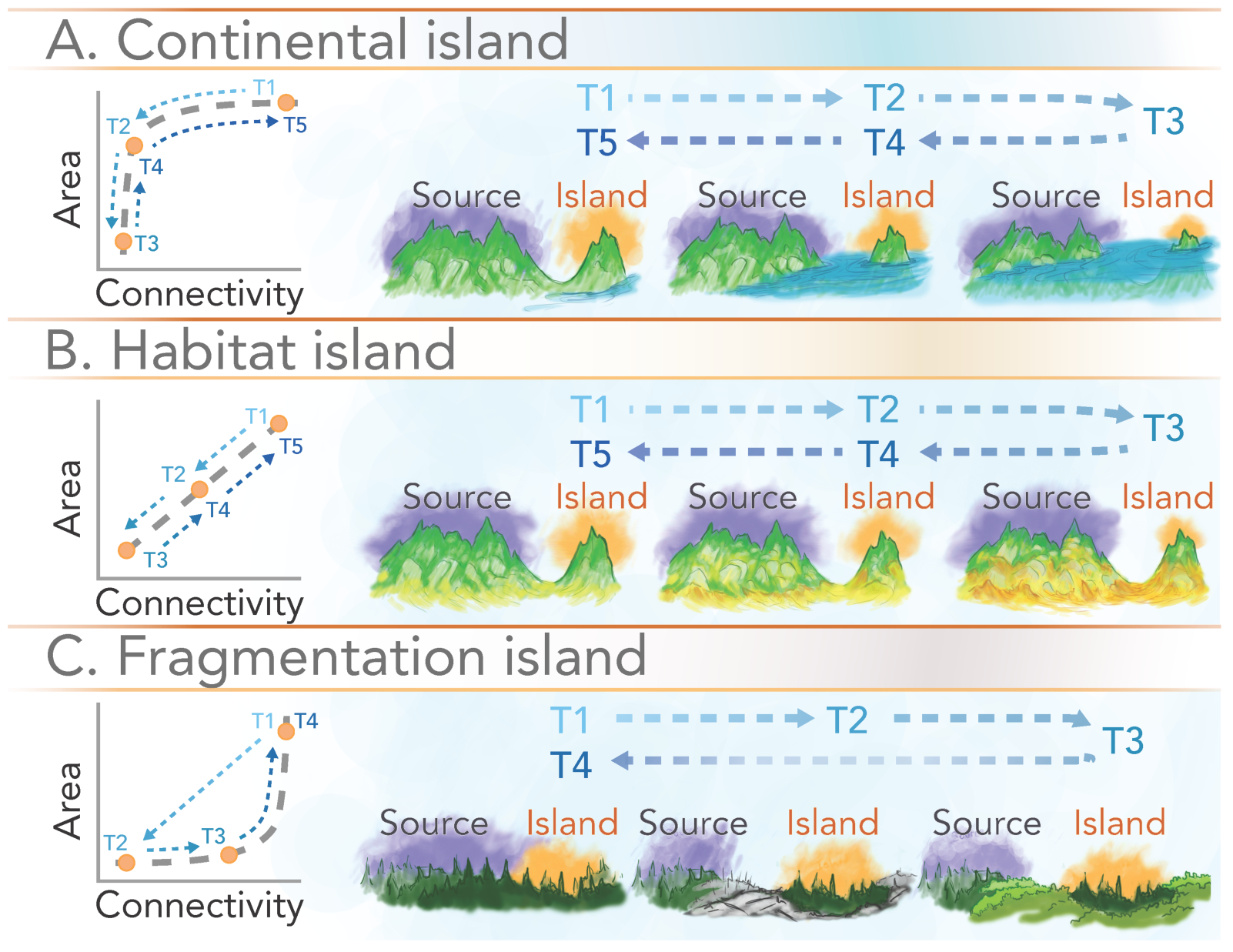

Types of island systems that undergo cyclical environmental changes and the predicted dynamic processes that drive candidate species pools on islands. Phases of colonization and extinction are not absolute, but represent periods dominated by the respective process. Oceanic islands most closely match Fragmentation islands that experience directional environmental changes, but not repeated cycles.

a) Continental or land bridge islands: Environmentally driven cyclical changes in sea level influence area and connectivity, with abrupt changes at critical thresholds of sea level. Islands form as they are isolated by rising sea level, followed by a gradual period of extinction as the island progressively decreases in size towards inundation. During the following period of recovery, connectivity will change little until sea level lowers enough to reestablish a connection with the mainland, prompting a rapid period of colonization.

b) Habitat islands (e.g., sky islands or other refugia of isolated habitat types): Long-term climatic cycles (e.g. glacial–interglacial) cause gradual changes in both area and connectivity. Gradual phases of extinction and colonization alternate, tracking environmental change. The system cycles repeatedly in a predictable way, although the duration and amplitude of each phase may vary.

c) Fragmentation islands: Over short time scales, natural or anthropogenic disturbances isolate habitat patches, which may eventually reconnect to contiguous areas of similar habitat as succession occurs within the intervening matrix. An initial wave of extinction accompanies a rapid decrease in area and connectivity. If recovery of the matrix is allowed to take place, a slower period of recolonization will occur as area and connectivity gradually increase. In different systems, cycles may occur repeatedly, with varying duration and predictability depending on the cause of disturbance.

Islands and habitat patches (e.g., insular systems) change in both size and connectivity to larger contiguous areas of similar habitat (e.g., mainland) over environmental cycles. Consequently, the composition of insular communities is driven by two distinct phases: i) colonization will lead to higher species richness during periods of increasing connectivity as the intervening matrix separating island from mainland becomes narrower and/or easier to traverse; and ii) extinction will reduce species richness during periods of decreasing area as an island’s capacity to support populations declines. The importance of these two processes was central to MacArthur & Wilson’s (1963, 1967) theory of island biogeography (TIB) which made predictions for species equilibrium richness on static islands with dynamic immigration and >extinction of species. Recent efforts have included the dynamic nature of insular systems (Heaney 2000; Heaney 2007; Whittaker et al. 2008; Warren et al. 2015; Patino et al. 2017; Whittaker et al. 2017). In parallel, a surge of studies has begun to incorporate organismal trait‐based approaches for studying ecology and biodiversity (e.g., McGill et al. 2006; Violle et al. 2014; Enquist et al. 2015). How trait-based constraints influence distributions on dynamic insular systems, however, remains poorly understood.

To address this shortcoming, we develop a model of how dynamic islands interact with biological constraints to influence colonization and extinction processes through environmental cycles. While the classic TIB predicted species richness (but not composition), recent extensions have begun to identify how differences in key functional traits can mediate colonization and extinction processes (Kirmer et al. 2008; Laurance 2008; Okie & Brown 2009; Jacquet et al. 2017). Thus, to predict populations and communities in insular systems, it is necessary to incorporate the interactions between organismal traits with the dynamic environmental constraints that drive alternating phases of colonization and extinction. Such considerations require explicit integration of the role of historical contingency or *hysteresis*—the dependency of outcomes not only on the current system state, but also the history of environmental change. Hysteresis has been identified as a property of many complex systems (Scheffer et al. 2001; May et al. 2008), and leveraging historical trajectories of insular systems may provide a powerful way to understand how species-level traits emerge as populations and communities (Chase 2003; Fukami & Nakajima 2011).

Accordingly, we present a Constraint-based model of Dynamic Island Biogeography (C-DIB) that integrates the cyclical nature of island size and connectivity with trait-based constraints on species to predict colonization and extinction probabilities of populations and communities through time. We present the two components of the model and then explore emergent properties at the population and community levels that are predicted. We begin with a simple framework that considers how species traits mediate demographic processes of populations in dynamic landscapes. Next, we explore how the population-level responses of species collectively form candidate pools for local communities. In the C-DIB model, colonization probability is constrained by traits that influence the dispersal capacity (e.g., mobility and physiological tolerances) of species to cross the intervening environmental matrix between island and mainland to establish a population. In contrast, extinction probability is driven by species traits (e.g., body size and trophic level) that constrain the minimum resource requirements (and hence area) necessary to maintain a viable population. Critically, because different suites of traits mediate the processes of colonization and extinction, the model reveals how identical system states — combinations of island connectivity and size — can result in different candidate species pools despite similar predicted richness. Our C-DIB model applies generally to a diversity of taxa and to insular systems that cycle on timescales of years and decades—such as disturbance and restoration—to tens of thousands of years—in the case of mountaintop and continental islands over glacial cycles (Box 1).

These advancements are timely for several reasons. First, anthropogenic environmental change is altering insular systems and creating new ones. Increasing temperatures cause mountaintop “island” habitats to contract (McDonald & Brown 1992; Dirnböck et al. 2011), while both continental and oceanic islands shrink until inundated from rising sea levels (Rahmstorf 2007; Whittaker et al. 2008; Weigelt et al. 2016). New islands are created as higher coastal areas become isolated. At the same time, local to regional anthropogenic disturbances across the earth lead to pervasive patterns of habitat fragmentation (Fahrig 2003; Laurance 2008; Haddad et al. 2015; Crooks et al. 2017). Efforts to predict changes in species distributions and anticipate extinctions are critical for effective conservation and restoration of biodiversity on a rapidly changing planet (McDonald & Brown 1992; Crooks 2002; Hampe & Petit 2005).

Second, by considering species trait-based constraints in a spatiotemporal environmental context, the model bridges key gaps in ecological theory by linking physical forces to population‐ and community-level dynamics (e.g., Pulliam 1988; Leibold et al. 2004; Agrawal et al. 2007). Specifically, it provides a predictive framework for integrating: i) trait-based constraints into landscape ecology, biogeography, and paleoecology (Jackson et al. 2009; Manning et al. 2009); ii) the effects of historical environmental trajectory into the same three subfields; iii) physiological constraints of individuals into population and community-level processes (McGill et al. 2006; Fristoe 2015); and iv) the effects of regional processes that filter local communities (Harrison & Cornell 2008).

Third, recent advances in theory, computational methods, and data from various fields now constitute a powerful toolkit to further understand dynamic populations and communities on islands and habitat patches. Metabolic scaling theory of resource allocation (e.g., Brown et al. 2004) provides a trait-based approach (e.g., Enquist et al. 2015) to understanding the role of body size, temperature, and trophic position on resource requirements, with predictable ramifications for life history, demography, and population dynamics. The compilation of large datasets of cross-species traits (e.g., PanTHERIA for mammals; Jones et al. 2009, BIEN for plants; Engemann et al. 2016) has allowed parameter estimations of scaling allometries that can lead to broad predictions, including population-level resource requirements and growth rates. The accumulation of rich data sources has also allowed key advances in methodologies of spatial mapping required to test the C-DIB model empirically. These include species occurrences (e.g., VertNet; Constable et al. 2010) and climate (e.g., WorldClim; Hijmans et al. 2005) used in developing Ecological Niche Models (a.k.a. Species Distribution Models but hereafter ENMs; e.g., Peterson et al. 2011). By using these tools and rich data-sources, we can begin to test predictions of the C-DIB model using contemporary systems, past systems using reconstructed paleo-climates, and future systems by forecasting anthropogenic change.

## A CONSTRAINT-BASED MODEL OF DYNAMIC ISLAND BIOGEOGRAPHY

We combine the processes of constraint-mediated colonization and extinction on dynamic island systems in a simple model. We begin with two processes well-established in the literature: trait-based colonization and extinction. We then build on these organismal constraints to show how their interaction with dynamic insular environments leads to novel predictions of hysteresis in island biogeography. The model predicts the presence or absence of populations of a given species at a particular time through environmental cycles (Box 2: Single species model). Additionally, when applied across all species present in the mainland source pool, the model predicts an island’s candidate species pool – a set of species available for local community assembly (Box 3: Community model). The model highlights the important role of hysteresis in affecting both species distributions as well as community composition, as past environmental states of the system affect the current presence or absence of species. The goal here is to present a simple and generalizable model with few assumptions of time lags, biotic interactions, and evolutionary stasis. We discuss later how these factors may be incorporated if specific systems require.

### Colonization

For a species to occur on a real or habitat island, individuals must first colonize from a source (mainland or nearby island) or have existed there before the island became isolated. Classically, non-random colonization ability has been defined through traits related to individual dispersal ability (Carlquist 1966; MacArthur & Wilson 1967). The processes of flying, rafting, or drifting correspond to quantifiable traits such as wing loading in birds (Hamilton 1961), swimming ability in terrestrial mammals (Meijaard 2001), and cluster size in seeds (Seidler & Plotkin 2006). Indeed, these traits facilitate crossing large habitat barriers. Similarly, some traits such as body size and home range constitute known predictors of dispersal distance in mammals, wherein larger species or those at higher trophic levels tend to have greater dispersal abilities, and therefore higher colonization potential (Bowman et al. 2002). Additionally, behaviors such as willingness to traverse open areas have been shown to affect colonization and distributions in insular systems (Moore et al. 2008).

In contrast, for systems where the distance between patches exceeds the dispersal ability of individual organisms, the primary driver of colonization probability shifts from individual dispersal to the establishment of populations across a matrix including corridors and steppingstones (Baum et al. 2004). At one extreme, the matrix separating islands from mainland can be entirely inhospitable. For example, the saline ocean that separates continental-shelf islands is physiologically intolerable to most amphibians, restricting colonization by mainland populations. However, this is not the case in many insular systems, where the peripheral habitats separating fragments from nearby areas of contiguous habitat are less discrete (Ricketts 2001; Laurance 2008; Prevedello & Vieira 2010). These matrices can be viewed as semi-permeable with suitability varying among species depending on their capacity for dispersal or population growth in conditions throughout the matrix. In these cases, contiguous or linked populations of some but not all species in the mainland source pool may become established on the island via corridors and stepping stones of suitable environments in a non-random way. In general, connectivity can be quantified as the environmental similarity of conditions in the matrix compared with those found on the island, with high connectivity relating to higher overall probabilities of colonization (Åberg et al. 1995; McRae et al. 2008; Altermatt et al. 2011; Holyoak 2014). Ecological niche modeling can be used to estimate species-specific environmental tolerances, allowing measures of matrix suitability (Peterson et al. 2011; Soley-Guardia et al. 2016) that indicate how colonization probabilities vary among species (Collinge 2000; Urban & Keitt 2001; Hunter 2002).

### Extinction

Local extinction or extirpation is driven by the effect of island area on the capacity of a species to maintain viable populations. In general, because larger islands can support a greater number of individuals, the probability of extinction will be lower than on smaller islands. Hence, larger islands show higher species richness (MacArthur & Wilson 1963, 1967). As an island shrinks due to environmental change, so does its capacity to support individuals of a given species, until it no longer provides sufficient area to maintain a viable population. This results in a non-random extinction process on islands due to variation in the spatial requirements of different species, which is mediated by organismal traits. Indeed, the literature documents several traits associated with differential extinction risk among species on islands including body size, trophic level, and degree of diet or habitat specialization (Brown 1971; Patterson 1987; Lessa & Farina 1996; Bolger et al. 1997; Larsen et al. 2005; Okie & Brown 2009; Boyer & Jetz 2014). These key traits influence the area necessary to support a minimum viable population (*sensu* Shaffer 1981; Soulé 1987) via individual resource requirements, space use, and population density.

Mammals provide a well-studied example. Larger species generally have greater per-capita energy requirements than smaller ones and population density scales negatively with body size (Damuth 1987; but see Silva & Downing 1995). Thus, larger islands with greater resource availability are necessary to sustain isolated populations of larger-bodied species (Marquet & Taper 1998). Likewise, mammals that feed at higher trophic levels have lower densities for a given body size (Carbone & Gittleman 2002; Marquet 2002) and hence cannot persist on small, isolated patches (Oksanen et al. 1981; Crooks 2002; Holt 2009). Finally, larger islands tend to contain a greater diversity of habitats and resource types, allowing more specialists to occur, whereas smaller islands should feature more generalists (Brown 1971).

### Predicting populations of a single species

In insular systems with dynamic conditions, the organismal traits mentioned above interact with changing environments to affect the presence or absence of a species. Because different suites of traits typically mediate the likelihood of successful colonization and the avoidance of extinction, the presence of a population of a particular species on a given island depends not only on current conditions, but also the history of the system. A population inhabiting an island will become extirpated if environmental change reduces the area of suitable habitat below the minimum threshold necessary to support a viable number of individuals (Keymer et al. 2000; Stephens 2016). Even if future environmental change restores conditions to a sufficiently large suitable area, a population will only reestablish if matrix conditions (i.e., distance or matrix composition) become amenable for colonization. Hence, due to hysteresis in the system, a species can be absent from an island it previously inhabited even when current conditions are favorable for supporting a population (e.g. Box 2, time T4, islands c and d). The presence or absence of a species is contingent on trajectories through previous environmental cycles. A species is only expected to occur in patches that have experienced a sufficient period of conditions favorable for colonization followed by a continuous period where island area is maintained above the minimum required to support a viable population (e.g. Box 2, T4, island b). As discussed earlier, species-specific traits affecting individual dispersal and population connectivity (e.g., Moore et al. 2008; Lasky et al. 2013) and those related to population maintenance (e.g., Brown 1971; Patterson 1987; Leibold et al. 2004; Okie & Brown 2009) mediate these respective probabilities of colonization and extinction.

#### Box 2.

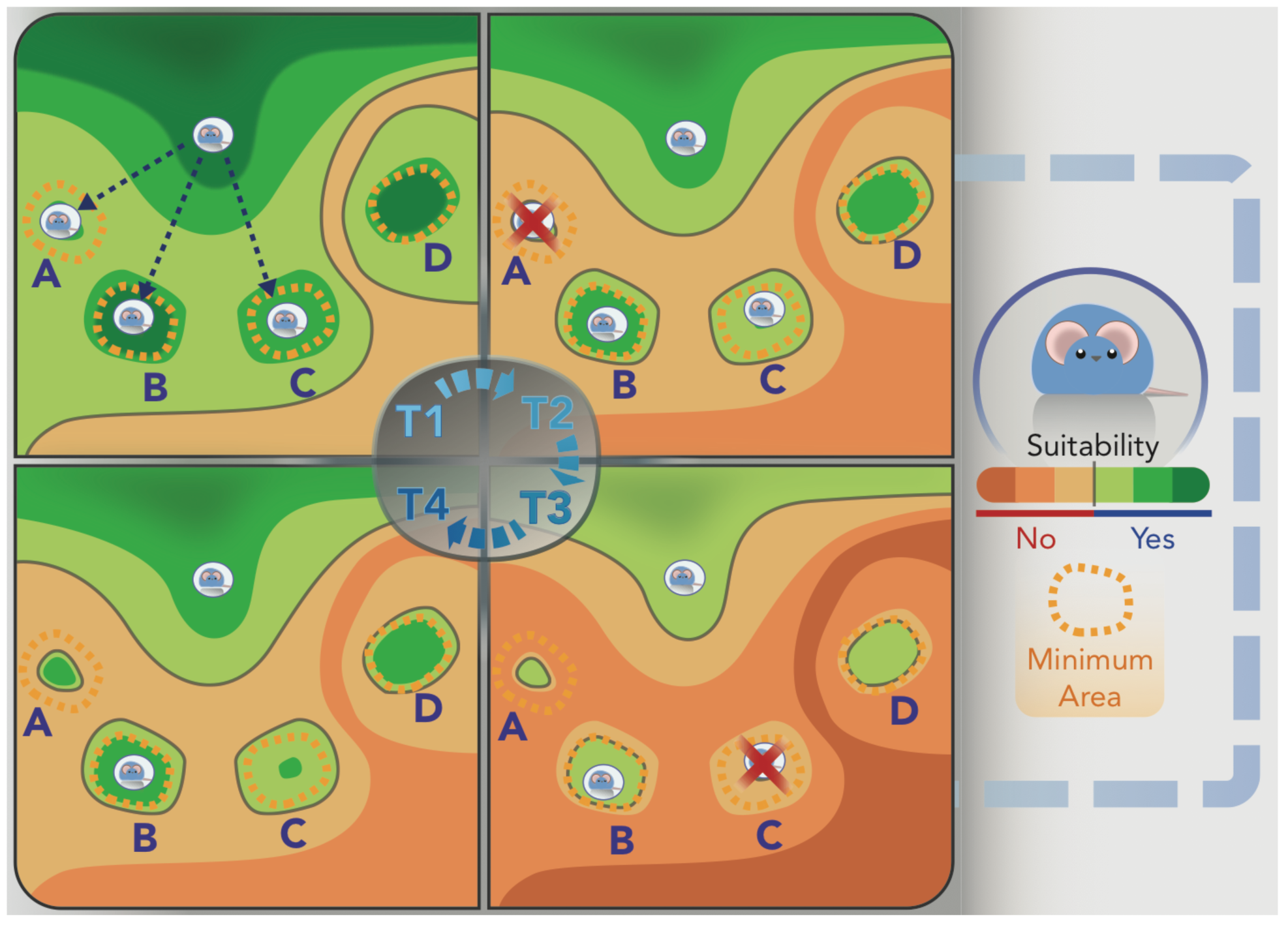

Single species model. The implications of the Constraint-based Dynamic Island Biogeography model are illustrated for populations of a single hypothetical species tied to mesic forest environments. Here, we consider a series of isolated mountaintop habitat islands of mesic forest surrounded by arid, non-forested areas as they are altered by climactic shifts associated with cycles of glaciation. During cooler, wetter glacial periods (T1), the forest habitat islands will be large, with the low valleys separating them from the ‘mainland’ (in this case a large, nearby mountain range) relatively mesic. This allows populations of our species to colonize islands a, b, and c. As the climate warms during an interglacial period, the area of suitable habitat will contract as cool, wetter climates and associated forests recede to higher elevations, while the intervening valleys become warmer, more arid, and less suitable (T1 –> T3). Over this period, populations will become extirpated from islands that drop below the minimum area necessary to support a viable number of individuals (islands a and c). Later as the environment rebounds (T3 -> T4), some islands will regain conditions favorable for supporting populations; however, without sufficient matrix suitability, they will not yet recolonize (T4, islands a and c). At time T4, the hypothetical species is only expected to occur on one island, where a period of conditions suitable for colonization were followed by continuous periods of island size above the minimum necessary to support a viable population (island b). Times T2 and T4 in the system demonstrate the influence of hysteresis, where our species is distributed differently across the islands, despite identical environmental conditions.

### Predicting candidate species for local community assembly

The combined processes of colonization and extinction of populations of each of the species present in the mainland pool predict the candidate species available for local insular communities. The model predicts asymmetrical changes in community composition for a given island over the course of environmental cycles (Box 3: community model). This means that particular species lost to extinction as island size and connectivity decrease, will not necessarily be the same species that recolonize as the island system reverses its trajectory and returns to earlier environmental states. Thus, an island can exhibit identical area and level of connectivity to the mainland at different times, but host a different set of species (e.g. Box 3, times T2a, T4a, T2b, and T6b, or times T3a, T3b, and T5b). This prediction bars any changes in the source pool on the mainland or intervening islands. Such discrepancies in composition at the community level is a consequence of hysteresis in the system. In operational terms candidate species will depend on: i) the previous island state and community composition, ii) the trajectory of the current cycle (see Box 3, environmental cycle 1), and iii) the magnitude of environmental change during previous cycles (see Box 3, environmental cycle 2).

#### Box 3.

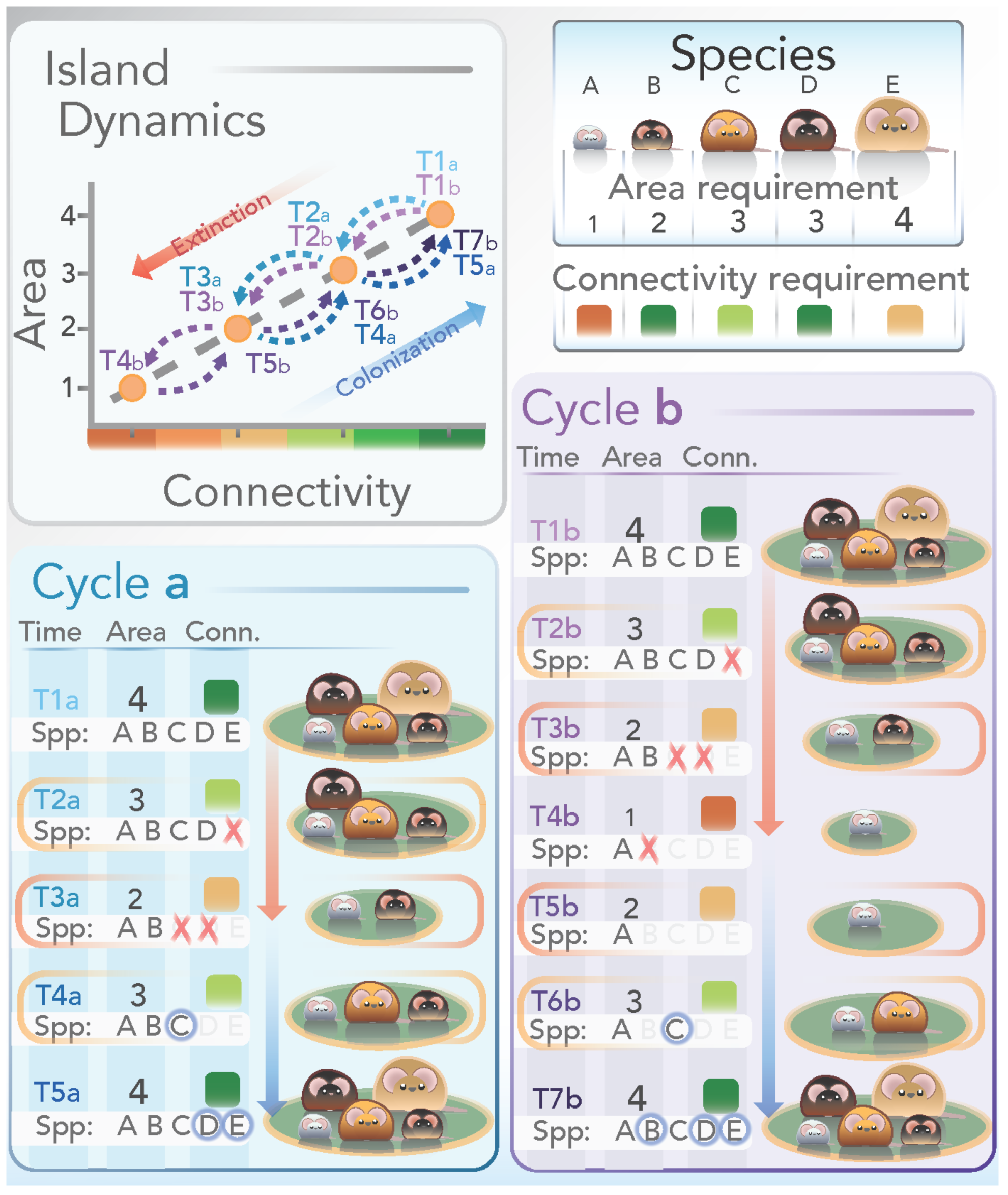

Community assembly model. To illustrate the consequences of the Constraint-based model of Dynamic Island Biogeography on community composition, we can consider a single hypothetical isolated mountaintop for a suite of species associated (to different degrees) with mesic forest. As the environment is altered by climatic shifts associated with glacial cycles, this mountaintop is periodically isolated by arid, non-forested areas. This example starts during a period of high connectivity when the island is inhabited by the full set of forest-restricted species found on a larger, nearby ‘mainland’ mountain range.

During these cool, wet glacial periods (e.g., T1a), the forest habitat island will be large and the low valleys that separate it from the ‘mainland’ range will be relatively mesic. As the climate warms during an interglacial period, the forest habitat will contract as moist climates recede to higher elevations while the intervening valleys become warmer and more arid (e.g. T1a -> T3a). Over this period, species will be lost to extinction as the island’s size decreases. The order that particular species disappear will be determined by traits influencing the area required for sustaining a viable population; in this case, we consider body size, with larger species requiring greater island area. Later, during the transition back towards a glacial period, as the climate cools and the island once again increases in size, so too will its capacity to support species (T3a -> T5a). However, a species’ likelihood of recolonization will depend on its ability to traverse the intervening matrix between island and mainland. In the case of the forest-dwelling animals considered here, the capacity to establish across the low-lying valleys at a given time is determined by tolerance to arid conditions (i.e. connectivity requirements). Following through environmental cycle a, demonstrates the role of hysteresis in structuring community composition as similar island conditions (the intermediate climates at T2a and T4a) are associated with different species. Not only the current position (i.e., environmental state), but also the current direction of the glacial cycle (trajectory in the cycle) has influenced the composition of species at this point in time.

The added influence of the magnitude of past environmental change is demonstrated by following the changes encountered over environmental cycle b. As the system returns to intermediate conditions (T6b) after experiencing a particularly warm interglacial period (T4b), the species present on the island differs from any other time with the same conditions that occurs during either environmental cycle a or b. Considering both cycles, bright lettering indicate time points that exhibit hysteresis, where identical island conditions are characterized by different communities (orange: times T2a, T4a, T2b, and T6b; and red: times T3a, T3b, and T5b).

## IMPLEMENTING THE MODEL

Useful functional traits for predicting colonization or extinction will vary across taxa and systems (e.g., Lenzner et al. 2017). The kind of island system (Box 1), as well as position and trajectory in the cycle, lead to testable predictions regarding the presence or absence of species’ populations and communities on islands through time. Active fields including ENM (e.g., Elith & Leathwick 2009) and metabolic scaling approaches (e.g., Sibly et al. 2012) provide practical tools for implementing the model across taxa. We present examples from a variety of insular systems that support components of the model to illustrate the broad utility of the C-DIB model.

### Model generality, trait selection, and island types

The C-DIB model applies generally to a wide range of taxa, island systems, and length of environmental cycle. Key, however, is that substantial trait variation exists among species resulting in variation in predictive value. Identifying useful traits requires a basic understanding of the system and taxa involved. For example, the conditions separating island from mainland can relate to a wide range of environmental filters—including gradients in temperature, soil acidity, vegetation type, ocean depth, or degree of fragmentation (Prevedello & Vieira 2010). In each case, different traits (e.g. thermal physiology, pH tolerance, soil and habitat preference) will interact with the environment to mediate colonization likelihoods and hence restrict species distributions. As suggested by McGill et al. (2006), for a trait to be useful in a comparative context, there should be greater among‐ rather than within-species variation by orders of magnitude.

While all insular systems are subject to the general predictions regarding colonization and extinction outlined in the previous sections, the dynamics of these processes will vary depending on the nature of environmental change (Box 1). Differences in island type will determine the duration and importance of extinction and colonization phases in driving population and community-level processes. In the case of continental islands, for example, extinction will be the dominant process until a connection with the mainland is reestablished, allowing species to recolonize (Box 1a). In contrast, in disturbance systems an initial wave of extinction will follow island formation followed by a period of gradual recolonization as the (semi-permeable) matrix recovers (Box 1c; Butaye et al. 2001). Such commonalities in island dynamics will influence the traits that are important for predicting species occurrences in particular systems. Some systems, taxa, and localities will undoubtedly reveal idiosyncrasies.

### Single versus multi-species comparisons

Tests of the model may be conducted at two primary levels as discussed above: single‐ and multi-species. Single-species (population-level) tests require trait data to calculate the probability of colonization and/or extinction. Importantly, it is not necessary to have complete trait information for the species studied. Researchers can leverage existing data and allometric scaling relationships to estimate missing ecological and life history parameters (see below). At the community level, however, more flexibility exists. Once again, species-specific trait data can be used to estimate colonization and extinction probabilities for each species in the mainland pool, leading to predicted candidate species for each island through time. Perhaps counterintuitive, community-level predictions can also be developed with less information. In cases where data remain insufficient for determining species-specific probabilities, trait information can be used to rank species in their order of colonization and extinction likelihood (Wright & Reeves 1992). This ranking approach allows tests of predicted community composition expected during colonization or extinction depending on the phase of the cycle.

## SPECIES DISTRIBUTION MODELS FOR ESTIMATING ENVIRONMENTAL SUITABILITY

Recent advances in modeling species niches and geographic distributions can be used to evaluate the C-DIB model using real and habitat islands. The difficulty in quantifying environmental suitability leading to estimates of connectivity among island patches has represented a major limiting factor in past research using traits to study the drivers of nested patterns of species distributions (e.g., Lomolino 1993). Ecological niche models quantify environmental suitability for a species based on occurrence records and environmental variables. Assuming niche conservatism (e.g., Wiens et al. 2010; Anderson 2013), ENMs can also be applied to reconstructed past or projected future conditions, permitting tests of the model through cycles over deeper time scales (Fig 1.). The field of ENM now includes options for supervised automation of advanced methods (e.g., via R, GUI, or other interfaces; Brown 2014; Hijmans et al. 2015; Wallace https://wallaceecomod.github.io) to facilitate tests of the model across many systems, taxa, and scales.

**Figure 1.**
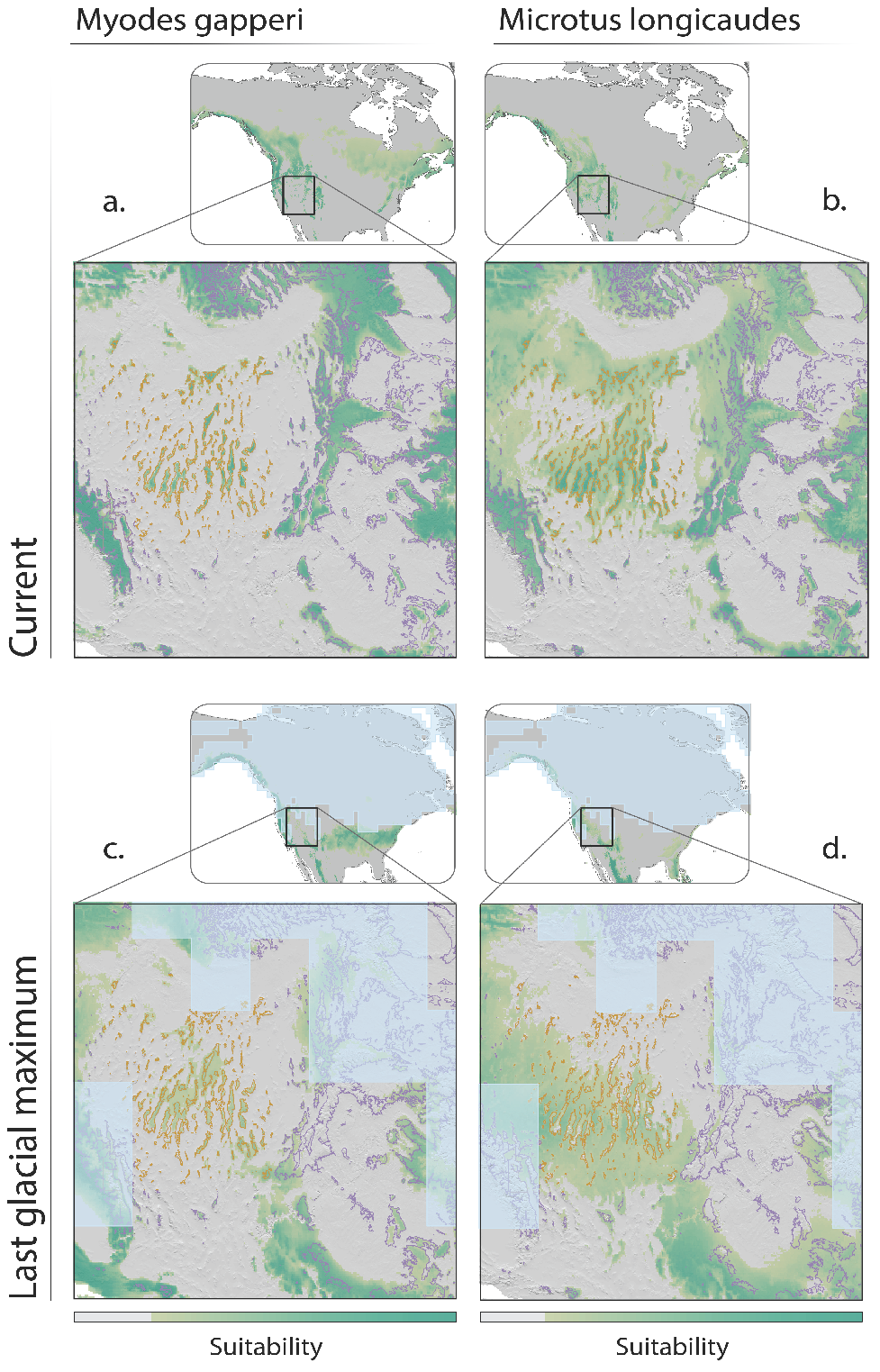
Colonization plot. Ecological niche models provide a valuable tool for quantifying differences in species-specific island connectivity currently and in the past. In analyses for two small mammals (voles), both *Myodes gapperi* (left) and *Microtus longicaudus* (right) are predicted to have suitable environmental conditions (darker green) at the present (top) on the larger sky islands of the Great Basin (elevations above 2200 meters outlined in orange). However, projecting those models to the last glacial maximum (bottom) indicates that only *M. longicaudus* had more extensive connectivity throughout the region across low-lying areas while *M. gapperi* has little. Likely source areas (i.e., the ‘mainland’ outlined in purple) occur in the Rocky Mountains (right side of maps) but also areas of the Sierra Nevadas (lower left portions of maps). This result is consistent with the current absence of *M. gapperi* from the Great Basin sky islands. The extent of glaciation during the last glacial maximum is depicted in blue (ENM estimates based on n = 245 and n = 309 point localities for *M. gapperi* and *M. longicaudus*, respectively; see supplemental).

Niche models can be implemented in various ways to evaluate different components of the C-DIB model. Such models can predict which species are likely to traverse the intervening matrix and colonize a patch. Habitat islands can be especially useful. First, delineation of suitable habitat is required to define the ‘island’, for example, by using a binary map of suitable/unsuitable to distinguish island from matrix at present, sets the stage for further analyses. This allows researchers to calculate the distance to an island, which is relevant for testing the C-DIB model using traits related to individual dispersal abilities. Furthermore, when applied to reconstructed past conditions, ENMs made for different species allow prediction of which species experienced contiguous corridors of suitable habitat connecting the mainland to the area that is presently an ‘island’ (e.g., at the Last Glacial Maximum; Waltari & Guralnick 2009).

The continuous quantification of suitability from such models reveals environmental gradients and can be combined with demographic analyses of population expansion (e.g., Anderson 2013) to complement research in landscape genetics, phylogeography, invasion biology, and estimates of range shifts under future climate change (Waltari & Guralnick 2009; Anderson et al. 2010; Prates et al. 2016). With additional information regarding population growth rates and dispersal kernels, such projections can estimate the colonization probability of populations over longer time scales. In addition to filling the void for estimating colonization probability across gradients, they also allow characterization of the size and environmental quality of each island, which is critical for estimating extinction probability (see below). These tools can refine our understanding of how populations and communities track environmental conditions as islands cycle through time.

### Modeling historical distributions across glacial cycles

Glacial-interglacial cycles serve as exemplary natural experiments to evaluate predictions of the C-DIB model over long time scales. Warming climate and rising sea level since the last glacial maximum continue to cause many mountaintop and continental shelf islands to shrink. It is not surprising that many such systems are currently undergoing community disassembly via the extinction process (Patterson 1987; Wright & Reeves 1992; Lomolino 1993; Okie & Brown 2009). The sky islands of the Great Basin have been instrumental in evaluating biogeographic theory by highlighting the role of historical factors that influence modern communities (e.g., Brown 1971; Rickart 2001; Rowe 2005, 2009). The extent of boreal forests in the Great Basin has cycled from expansive coverage including both mountains and low-lying areas during glacial maxima, to isolated patches on mountaintops surrounded by a matrix of lowland desert scrub during interglacials. These sky islands have been isolated from more contiguous boreal habitats in the Rocky Mountains since the earth’s shift out of the last glacial maximum, although the highest-elevation vegetation types (yellow pine, spruce, and fir) likely were not connected even during glacial times (Brown 1971). Waltari and Guralnick (2009) used ENMs and physiographic paleontological maps combining elevation, hydrology, and climate to quantify areas of suitable environment for montane mammals found on sky islands of the Great Basin both in the present and during the last glacial maximum. Important for testing the C-DIB model, their results show that habitat connectivity varies among species and between time periods. Furthermore, several species inhabiting nearby ranges of the Rocky Mountains, particularly those associated with yellow pine, spruce, and fir forests, are not found in the sky islands despite the current presence of suitable habitat there today (Brown 1971).

Figure 1 illustrates the utility of projecting historical distributions to understand variation in colonization probabilities. We contrast the estimated ENMs for two rodent species, *Microtus longicaudus* (currently found in the sky islands and the Rockies) and *Myodes gapperi* (currently restricted to the Rockies). Doing so indicates that there are currently suitable conditions throughout the sky islands for both species (Fig 1a-b). However, hindcasting the models to the last glacial maximum suggests that suitable areas for *M. gapperi* in the Rocky Mountains were not connected to those occurring in the sky islands (Fig 1c). On the other hand, suitable conditions for *M. longicaudus* spanned the entire region (including the lowlands), suggesting that past corridors facilitated colonization to the sky islands (Fig 1d). This provides a preliminary example of how ENMs can inform the colonization process of the C-DIB model by revealing variation in suitable corridors of connectivity among island patches, projected back during the last glacial maximum. These same methods can be applied to deeper time scales (e.g., Peterson & Ammann 2013), as well as coupled with population-genetic studies (Crespi et al. 2003; Floyd et al. 2005; Prates et al. 2016).

## IMPOSING ALLOMETRIC CONSTRAINTS

Ecological niche models can be combined with allometric constraints to improve predictions of an island’s capacity for sustaining populations of given species, providing a cutoff for islands that are too small and disconnected. For example, mechanistic approaches link ENMs to physiological processes that emerge at the level of populations and geographic distributions (e.g., Kearney & Porter 2009). Body size is an easily quantifiable trait (including from fossils), and many characteristics of organisms scale predictably with size such as population densities (Damuth 1981), home range (Kelt & Van Vuren 2001; Tamburello et al. 2015), and population growth rates (Sibly et al. 2012). In general, these allometric scaling relationships take power law form, *Y = Y*_*o*_*M*^*b*^, where *Y* is a trait of interest such as individual area requirements, *M* is body size, *Y*_*o*_ is the intercept, and *b* is the slope. Across large taxonomic scales spanning orders of magnitude in body size, individual resource requirements reflected in metabolic rates generally scale as *M*^3/4^ (Brown et al. 2004) and maximum population densities scale as *M*^-3/4^ (Damuth 1981). The result is that larger species generally occur at lower densities and therefore require larger areas to sustain the same number of individuals (Marquet & Taper 1998; Okie & Brown 2009).

In addition to body size, the elevation (i.e., intercept) of allometric scaling relationships provide a useful tool for evaluating the effects of trophic level on the process of the C-DIB model. Trophic position is a useful trait that is easily inferred from morphology, dentition, stomach contents, and stable isotope analysis. In mammals, the height of the coefficient (*Y*_*o*_) in density-size scaling relationships is influenced by trophic position, such that primary consumers occur at greater population densities than omnivores and carnivores after accounting for body size (Carbone & Gittleman 2002; Marquet 2002; Burger et al. 2017). This framework predicts that the minimum island size suitable for supporting an herbivore population would have to be at least ∼10 times larger to support a population of a similar sized carnivore (Fig 2). The scaling of ecological traits with body size and trophic level provides an operational framework to define the minimum area required to sustain a viable population.

**Figure 2.**
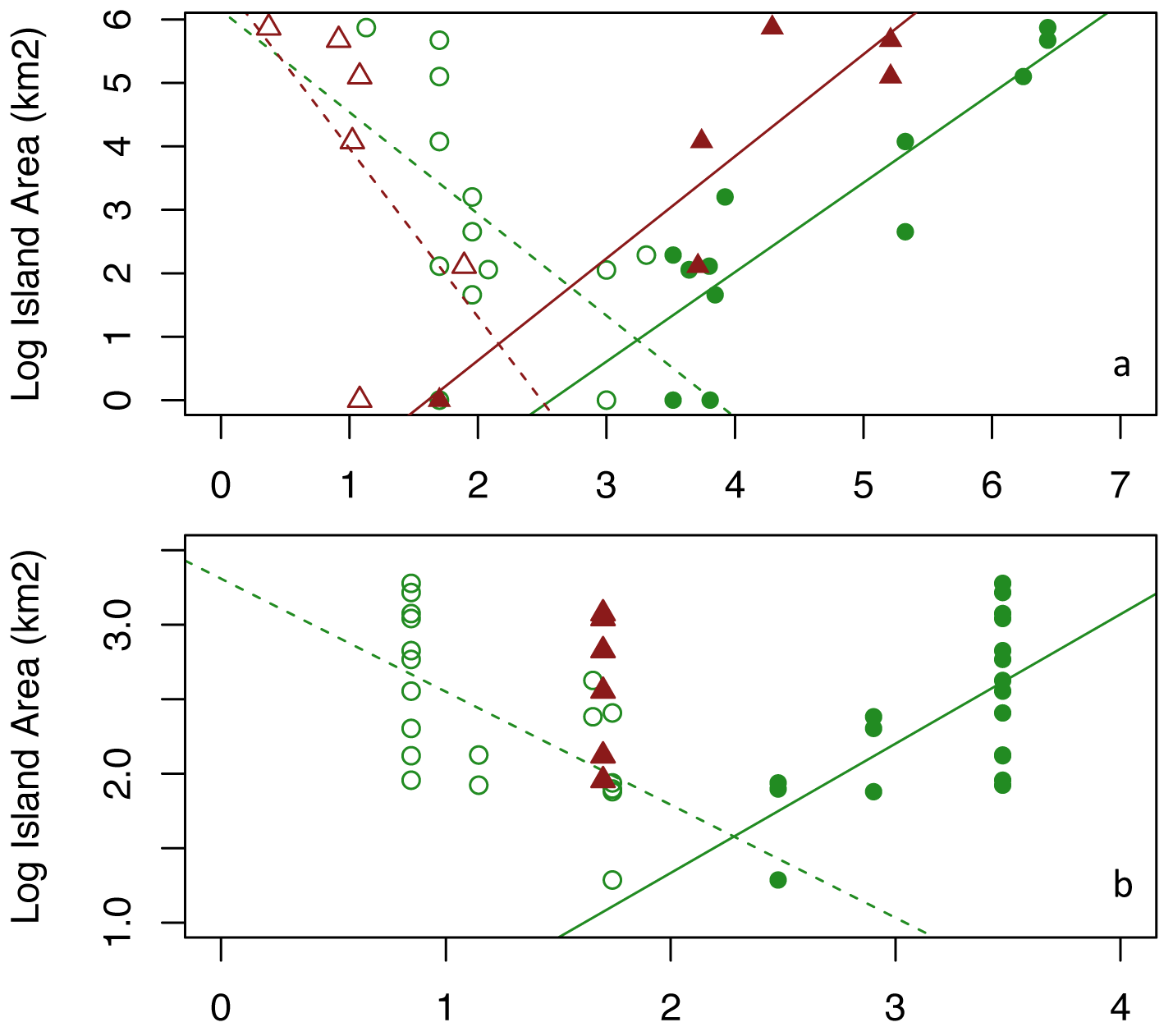
Extinction plots. Non-random assembly of body size and trophic distributions of mammals of: a) the Sunda Shelf continental islands of Indonesia and parts of Malaysia, and b) the great basin “sky island” mountaintops of the western USA. Open symbols = smallest body sizes and filled = largest body sizes. Primary consumers are green circles and secondary consumers are red triangles. Regression fits to largest (solid) and smallest (dashed) body sizes for each island system show significant convergence to a smaller range of body sizes on smaller islands in both insular systems. Note the left shift in body sizes for carnivores compared to herbivores supporting predictions of trophic energetics. Only one carnivore (*Mustela erminea cicognani*; 55 grams) appears on the largest GB “sky islands” and is absent from the smallest islands. Data for the Great Basin sky islands are from Brown (1978) and Grayson and Livingston (1993) and Sunda Shelf from Okie and Brown (2009).

Allometries can also inform estimates of demographic growth rates and colonization fronts. Mass-specific metabolic rate scales as *M*^-1/4^ in mammals, resulting in other biological rates such as reproduction and population growth rates (including *r max* or maximum intrinsic growth rate) to scale as *M*^-1/4^ (Blueweiss et al. 1978; Hennemann 1983; Calder 1984; Peters 1986; Sibly et al. 2012). Therefore, combining allometric constraints with ENMs discussed above can allow prediction of the maximum speed at which species can colonize through a formerly inhospitable intervening matrix that has recently become suitable, informing concerns for time lags (see below).

### Converging extinction processes on islands

While the simple power laws discussed above provide general rules to guide initial predictions, it should be noted that for certain taxa some ecological attributes, such as population density, may not scale monotonically with body size. For example, in mammals the highest growth rates and densities occur at intermediate body sizes near the metabolically optimal ∼100 grams (Silva & Downing 1995) as evident on islands (Marquet & Taper 1998). These multiple scaling relationships allow estimates of trait-based constraints on populations, which can be used to evaluate the extinction process of the C-DIB model even for species with missing data. This highlights the need to understand how constraints linked to different traits that can act on population colonization and extinction may vary depending on taxa (e.g., Jacquet et al. 2017).

The mammal faunas of two distinct insular systems – the continental islands of the Sunda Shelf and the ‘sky islands’ of the Great Basin of Western North America – reveal illuminating examples of the similar convergent patterns of non-random extinction due to the island size reduction process of the C-DIB model (Fig 2). Supporting predictions of trait-mediated extinction, the smallest patches in these island systems host mammalian communities characterized not only by lower richness, but also i) reduction in the range of body sizes present, ii) fewer species at high trophic levels, and iii) lack of specialists (Brown 1971; Weigelt et al. 2016; Okie & Brown 2009). These natural experiments of extinction patterns among the smallest and largest body sizes provide preliminary evidence that energetic constraints on sustaining small populations may be widely important and predictable. For large and small species, the number of individuals that can exist in a given area is low, consistent with extinction risk increasing as area shrinks. Each species has a minimum threshold area below which a minimum viable population is unable to persist.

## PREDICTING FUTURE POPULATIONS AND COMMUNITIES ON FRAGMENTS

In addition to implementing the model for understanding species responses to climate change over long time scales of 10,000-100,000 years, the C-DIB model also provides predictive power for how communities will respond in the near-term to habitat fragmentation caused by natural and anthropogenic habitat disturbances (e.g. wildfires or clearcut logging) on time scales of years to decades. Just as glacial cycles provide “natural experiments” over the long-term, human-induced disturbance and subsequent recovery provide “anthropogenic experiments” to test the model on much shorter time scales (e.g., Soulé et al. 1992). Despite variation by orders of magnitude in the temporal length of cycles, these kinds of systems are remarkably similar in their cyclical nature. In both cases, habitat patches cycle from larger and more connected patches to smaller, isolated fragments. If allowed to recover in the case of disturbances, this process is reversed and patches become larger and more connected (Box 1).

A key advantage of studying such fragmentation/succession island systems is that cycles occurring over short time scales provide opportunities to observe the process of island colonization (e.g., Simberloff & Wilson 1969), allowing experimental tests of C-DIB model predictions. If habitats within the matrix are allowed to recover following a disturbance, the colonization probability of species will generally rise as connectivity between island and nearby areas of contiguous habitat increases. Trait data on dispersal ability, physiological tolerances, or habitat associations allow predictions of the sequence in which species will re-establish. For example, as a previously forested (and then clear-cut) region recovers, species more tolerant of secondary growth would be expected to colonize expanding remnant habitat patches earlier than those that only occur in mature growth (Stouffer & Bierregaard 1995). Similarly, the C-DIB model can inform restoration efforts aiming to re-establish populations in previously disturbed habitat patches (e.g., McIntire et al. 2007; Cristofoli et al. 2010). In these cases, the area and quality of the habitat patch will increase gradually, and colonization will often take place across an unchanging intervening matrix. These systems will be characterized by a period of increasing richness as establishment of species occurs. The C-DIB model predicts which kinds of species will colonize and in which order.

Conversely, if a disturbance episode fragments a contiguous habitat into isolated patches, local populations are lost to trait-biased extinctions. “Anthropogenic experiments” in the form of habitat fragmentation in tropical forests provide instructive examples. For instance, the creation of a hydro-electric lake, the flooding of low-lying areas, and the subsequent islands formed in tropical forests of Venezuela led to the loss of large mammals at higher trophic levels (Terborgh et al. 1999)—a prediction of the C-DIB model. Similarly, the Biological Dynamics of Forest Fragments Project in Brazil (Bierregaard et al. 1992) provides another opportunity to apply the C-DIB model. As a planned experiment, forest plots varying in size (ranging from 1-100 ha) were left intact as surrounding forests were cleared. Researchers inventoried species composition in fragments before and after forest clearing, collecting data on a diversity of taxa including birds, plants, small mammals, primates, insects, and frogs (Laurance et al. 2002). Extirpations in forest fragments provide opportunities to evaluate the importance of different traits in determining extinction risk and to predict future species losses. Conversely, an experiment of successional-forests grids in northeastern Kansas allows examination of the opposing directional process. Old-field environments (long disturbed by agriculture) were allowed to regrow towards the woody climax community possible in that region (Diffendorfer et al. 1995; Holt et al. 1995); however, succession was only permitted in particular blocks that varied in size and distance to the nearest large patch of preexisting forest. These and other anthropogenic experiments can be used to evaluate predictions of the C-DIB model on short time scales.

## FROM CANDIDATE SPECIES POOLS TO REALIZED COMMUNITIES

While recognizing the inherent idiosyncrasies of many natural systems, we have presented a model that is deliberately simple with the aim of making general predictions using the fewest possible parameters (Marquet et al. 2014). By illustrating how key species traits interact with environmental constraints over time, the C-DIB model predicts a pool of candidate species available for island community assembly. As mentioned in the Introduction, a number of additional processes will subsequently filter or facilitate the predicted candidate species pool into the set of species that ultimately occupy any element of an island system at a particular point in time. We present a selection of potentially important processes that future work may integrate into the C-DIB framework.

### Time lags, extinction debts, and colonization credits

Extinction debts and colonization credits (Jackson & Sax 2010) are likely when the rate of environmental change is fast relative to the demographic rates of focal taxa. The C-DIB model makes predictions regarding how historical changes in environmental conditions influence the presence or absence of a particular species, assuming equilibrium with the environment for a given ‘snapshot’ in time. However, the processes of colonization and extinction are complicated due to: i) stochastic elements of patch occupancy (Hanski & Ovaskainen 2003), ii) relative rates in the changes in the environment, and iii) population dynamics of the species involved. Because colonization and extinction are probabilistic processes, time becomes an important factor in the likelihood of either event occurring. The longer island conditions remain below the minimum necessary to sustain a population of a given species, the more likely the species is lost from the insular community. This can result in time lags and discrepancies between theoretical predictions and empirical data (Jackson et al. 2009). When matrix conditions allow for colonization, it is possible for an individual of a given species to be present even if island conditions are not sufficient to support a self-sustaining population (i.e., island size is too small), as emigrating individuals from a source population on the mainland or another island maintain the island sink population (e.g., rescue effect; Brown & Kodric-Brown 1977).

The C-DIB model can be extended to make predictions about extinction debts—when a species is currently present but doomed to extinction because patch size has been reduced below the minimum threshold necessary to sustain a population (Keymer et al. 2000; Kitzes & Harte 2015). Conversely, in cases of habitat restoration, the model can inform colonization credits—where a species is not yet present, but restored patches include the necessary conditions for colonization and subsequent persistence. These realities will inevitably result in discrepancies between the predicted candidate species pool and realized community composition (Tilman et al. 1994; Cristofoli et al. 2010; Kitzes & Harte 2015). Such considerations also apply directly to the expansion and contraction of species distributions under dynamic environments, where population-level responses may or may not track environmental change (Engler et al. 2009; Anderson 2013; Estrada et al. 2015).

### Biotic interactions

The C-DIB model presented here also ignores the effects of biotic interactions that undoubtedly play crucial roles in shaping local populations and communities on islands (Vannette & Fukami 2014). The establishment and persistence of a particular species clearly impacts and is impacted by interactions with other species. These include positive and negative effects such as exclusion, facilitation, trophic release, and priority effects, among others (Cody & Diamond 1975; Fukami 2015). Despite the difficulty in detecting the effects of biotic interactions on species ranges and taking them into account in modeling their niches/distributions (Anderson 2017), recent work has made progress in elucidating the role of species traits in mediating direct and indirect interactions within populations and communities (Bolker et al. 2003; Werner & Peacor 2003; Lessard et al. 2016). An example of the potential implications regarding trait-mediated extinction is that as patches are reduced, predators become extirpated first, causing cascading effects that lead to increased herbivory as primary consumers exhaust their resources and exceed carrying capacity (e.g., Estes et al. 2011). Understanding which traits affect the presence and intensity of both negative and positive interactions has potential to expand the C-DIB model by further narrowing the candidate species pool, or widening it in the case of facilitation (Bruno et al. 2003). This can further complicate predicting metacommunities where species are linked by dispersal of multiple interacting species (Leibold et al. 2004). These issues may be resolved by considering the likelihood of subpopulations and sub-communities insufficient to sustain isolated populations (i.e., sinks; Pulliam 1988; Hanski 1999) by combining ENMs and allometric constraints discussed above.

### Evolution on islands

Finally, the C-DIB model does not currently account for the effects of evolutionary processes on insular systems. Indeed, speciation has been the focus of several dynamic island models (Heaney 2000; Whittaker et al. 2008; Steinbauer et al. 2013). However, speciation per se may not be of large consequence when predicting the past or future distribution of species, even over a full glacial–interglacial cycle (Heaney 2000). Speciation may be a larger factor in older and more distant islands (Whittaker et al. 2008). However, the power of evolutionary and cultural adaptations to shape relevant species traits raises issues of genetic drift and local adaptation (Leimu & Fischer 2008). While trait evolution is not as likely to be an issue when considering islands that cycle on relatively short time scales (e.g. disturbance-succession islands), it becomes important over increasing time scales (e.g., Steinbauer et al. 2012). Glacial cycles occur on time scales of 10,000-100,000 years (Milankovitch 1941; Berger 1988; Roy et al. 1996), whereas species duration, especially for vertebrates and vascular plants, corresponds to much longer time scales (Bennett 1990; Kidwell 2013). Therefore, assumptions of niche conservatism over glacial cycles (e.g., Waltari & Guralnick 2009) are likely met for large organisms with slow generation times. However, issues may arise for small organisms with fast generation times, such as microbes. System size and time scale of cycle relative to organismal size and generation times constitute fascinating and important issues that need consideration in different systems. System time-cycles and the dependency of generation time on body size and temperature as predicted by metabolic theory (Savage et al. 2004) can evaluate this assumption of the C-DIB model and lead to subsequent extensions of it.

## CODA

Our C-DIB model builds on foundations laid by MacArthur & Wilson (1963, 1967) by incorporating trait-based constraints on colonization and extinction of insular systems with environmental and spatial characteristics that cycle through time. Notably, the C-DIB model leads to novel predictions of hysteresis in island biogeography. The model has important implications for understanding the effects of rapid habitat fragmentation on biodiversity—a pervasive feature of the Earth’s landscapes and seascapes. Recent availability of rich datasets, trait-based theory, and computational tools showcase exciting opportunities to evaluate the CDIB model in various systems and taxa. Doing so will improve our understanding of the physical forces and biological constraints that act on populations and communities to make up the spectacular diversity of life on the planet.

## Acknowledgments

We thank members and affiliates of the Anderson lab at CUNY and Peter White and the Hurlbert lab at UNC for comments on the manuscript. James Brown, Jordan Okie, and Mariano Soley-Guardia provided helpful discussions. Bruno Vilela gave advice on ecological niche models. RPA acknowledges funding from the U.S. National Science Foundation (NSF DEB-1119915 and DBI-1650241). JRB was supported by a Carolina Postdoctoral Fellowship for Faculty Diversity. Our collaboration was catalyzed by the University of New Mexico’s Biology Seminar series in 2015.

---

**Supplemental Figure 1.**
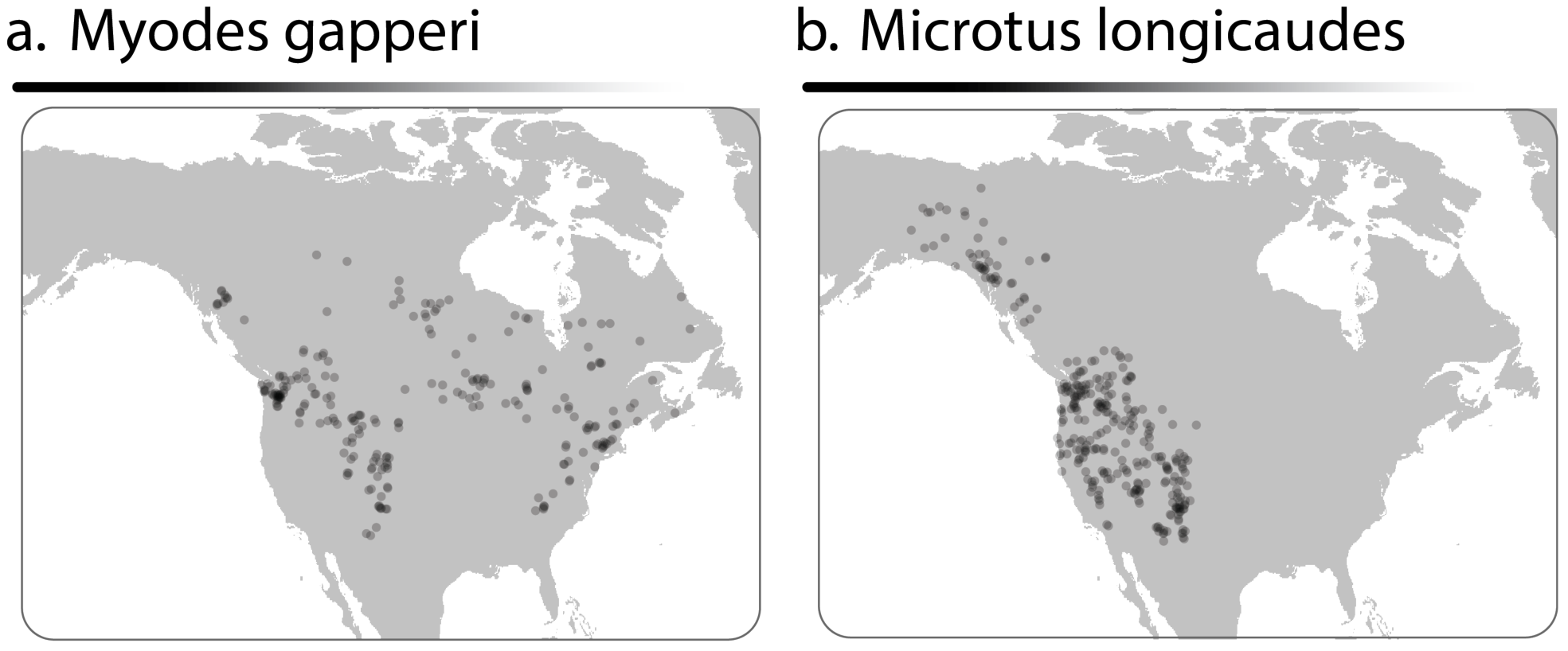
Point localities used in estimating ecological niche models for *Myodes gapperi* (a; n = 245) and *Microtus longicaudus* (b; n = 309).

**Supplemental methods for Figure 1.** Code for estimating ecological niche models was modified from R scripts originally produced in the Wallace package (version 0.6.4; https://cran.rproject.org/web/packages/wallace/index.html). For biological input data, we emulate data sources from Waltari & Guralnick (2009). For each species, 10,000 occurrence records were downloaded from the Global Biodiversity Information Facility (https://www.gbif.org), using only those with georeferences with duplicate locations removed. To increase reliability of species identifications, records were reduced to include only those from the Biodiversity Research Center at the University of Kansas, the Field Museum of Natural History, the Florida Museum of Natural History, Los Angeles County Museum, Louisiana State University Museum of Natural Science, Michigan State University Museum, Museum of Southwestern Biology, Royal Ontario Museum, Santa Barbara Museum of Natural History, Slater Museum of Natural history at University of Puget Sound, Sternberg Museum of Natural History at Fort Hayes, University of Alaska Museum of the North, University of California Berkeley Museum of Vertebrate Zoology, University of Colorado Museum of Natural History, University of Washington Thomas Burke Memorial Museum, and the Yale University Peabody Museum of Natural History. Records from outside of the species’ known distribution (range map downloaded form IUCN) were excluded from analyses. Remaining occurrence localities were spatially thinned to 10 km to reduce effects of biased sampling, yielding n = 245 localities for *M. gapperi* and n = 309 for *M. longicaudus*. Environmental data for the current and last glacial maximum included bioclimactic variables from WorldClim at 2.5arcmin resolution (www.worldclim.org). Ten thousand random background points were selected from within a minimum bounding polygon that included all localities and a 2 degree buffer. Model predictive performances were tested using a cross-validation approach that partitions localities into four spatially delimited blocks with an aggregation factor of 2. We made Maxent models of various complexities, specifically L, LQ, and LQH feature classes with regularization multipliers spanning from 1 to 5 by 0.5 intervals (L = linear; Q = quadratic; H = hinge). To approximate optimal model complexity, we chose the model with the lowest AICc (*M. gapperi*, supplemental table 1; *M. longicaudus*, supplemental table 2). Model outputs were visualized using the logistic transformation, and areas below the 10 percent threshold appear transparent (*M. gapperi*, 0.20; *M. longicaudus*, 0.16). The extent of glaciation during this time period is from CLIMAP (https://iridl.ldeo.columbia.edu/SOURCES/.CLIMAP/.LGM/index.html?Set-Language=en).

**Supplemental table 1.** Evaluation statistics for ecological niche models for *Miodes gapperi*.

| Feature classes | RM | Full AUC | Mean AUC test | Mean AUC Difference | Mean OR | AICc | Delta AICc | number parameters |
| --- | --- | --- | --- | --- | --- | --- | --- | --- |
| LQH | 5 | 0.85 | 0.75 | 0.16 | 0.34 | 6246.63 | 0.00 | 28.00 |
| LQH | 4.5 | 0.86 | 0.75 | 0.16 | 0.34 | 6249.50 | 2.87 | 32.00 |
| LQH | 3.5 | 0.86 | 0.77 | 0.14 | 0.33 | 6251.75 | 5.11 | 41.00 |
| LQH | 4 | 0.86 | 0.76 | 0.15 | 0.33 | 6259.38 | 12.74 | 39.00 |
| LQH | 3 | 0.86 | 0.76 | 0.15 | 0.34 | 6264.38 | 17.75 | 46.00 |
| LQ | 1.5 | 0.84 | 0.68 | 0.21 | 0.39 | 6277.71 | 31.08 | 23.00 |
| LQ | 1 | 0.84 | 0.70 | 0.20 | 0.38 | 6281.21 | 34.58 | 26.00 |
| LQ | 2.5 | 0.84 | 0.68 | 0.19 | 0.39 | 6281.54 | 34.90 | 24.00 |
| LQ | 2 | 0.84 | 0.68 | 0.20 | 0.39 | 6283.81 | 37.17 | 26.00 |
| LQ | 3.5 | 0.84 | 0.70 | 0.19 | 0.39 | 6286.65 | 40.01 | 24.00 |
| LQ | 3 | 0.84 | 0.70 | 0.19 | 0.38 | 6288.19 | 41.56 | 24.00 |
| LQ | 4 | 0.84 | 0.70 | 0.19 | 0.39 | 6294.70 | 48.07 | 25.00 |
| LQ | 4.5 | 0.84 | 0.70 | 0.19 | 0.39 | 6303.33 | 56.69 | 24.00 |
| LQ | 5 | 0.83 | 0.71 | 0.19 | 0.39 | 6305.00 | 58.37 | 21.00 |
| L | 1 | 0.78 | 0.69 | 0.16 | 0.28 | 6401.64 | 155.01 | 13.00 |
| L | 1.5 | 0.78 | 0.70 | 0.16 | 0.27 | 6401.73 | 155.10 | 13.00 |
| L | 2 | 0.78 | 0.70 | 0.15 | 0.25 | 6408.16 | 161.52 | 14.00 |
| L | 4.5 | 0.78 | 0.72 | 0.13 | 0.24 | 6409.04 | 162.41 | 11.00 |
| L | 3.5 | 0.78 | 0.72 | 0.14 | 0.25 | 6409.31 | 162.68 | 12.00 |
| L | 5 | 0.78 | 0.72 | 0.13 | 0.23 | 6409.66 | 163.02 | 11.00 |
| L | 3 | 0.78 | 0.71 | 0.14 | 0.25 | 6409.93 | 163.30 | 13.00 |
| L | 2.5 | 0.78 | 0.71 | 0.14 | 0.26 | 6410.39 | 163.75 | 14.00 |
| L | 4 | 0.78 | 0.72 | 0.13 | 0.24 | 6410.45 | 163.81 | 12.00 |
| LQH | 2.5 | 0.87 | 0.75 | 0.16 | 0.35 | 6767.05 | 520.41 | 54.00 |
| LQH | 2 | 0.87 | 0.80 | 0.07 | 0.25 | 6977.87 | 731.23 | 45.00 |
| LQH | 1.5 | 0.86 | 0.75 | 0.12 | 0.34 | 7022.75 | 776.12 | 35.00 |
| LQH | 1 | 0.51 | 0.64 | 0.02 | 0.10 | NA | NA | 66.00 |

L = linear; Q = quadratic; H = hinge; RM = regularization multiplier; AUC = area under the curve of the receiver operator characteristic plot; OR = omission rate.

**Supplemental table 2.**
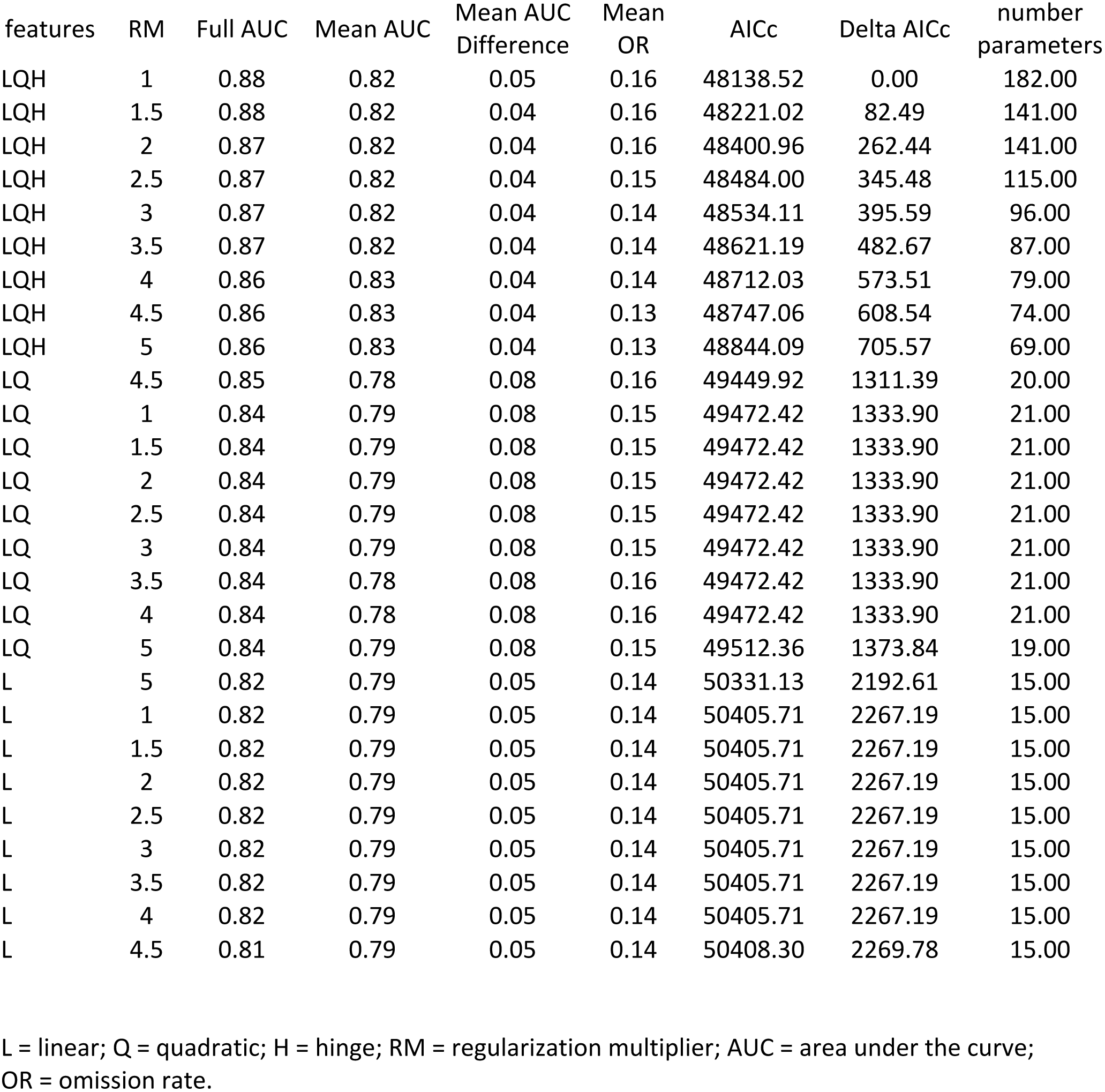
Evaluation statistics for ecological niche models for *Microtus laungicaudus*.

| features | RM | Full AUC | Mean AUC | Mean AUC<br>Difference | Mean<br>OR | AICc | Delta AICc | number<br>parameters |
| --- | --- | --- | --- | --- | --- | --- | --- | --- |
| LQH | 1 | 0.88 | 0.82 | 0.05 | 0.16 | 48138.52 | 0.00 | 182.00 |
| LQH | 1.5 | 0.88 | 0.82 | 0.04 | 0.16 | 48221.02 | 82.49 | 141.00 |
| LQH | 2 | 0.87 | 0.82 | 0.04 | 0.16 | 48400.96 | 262.44 | 141.00 |
| LQH | 2.5 | 0.87 | 0.82 | 0.04 | 0.15 | 48484.00 | 345.48 | 115.00 |
| LQH | 3 | 0.87 | 0.82 | 0.04 | 0.14 | 48534.11 | 395.59 | 96.00 |
| LQH | 3.5 | 0.87 | 0.82 | 0.04 | 0.14 | 48621.19 | 482.67 | 87.00 |
| LQH | 4 | 0.86 | 0.83 | 0.04 | 0.14 | 48712.03 | 573.51 | 79.00 |
| LQH | 4.5 | 0.86 | 0.83 | 0.04 | 0.13 | 48747.06 | 608.54 | 74.00 |
| LQH | 5 | 0.86 | 0.83 | 0.04 | 0.13 | 48844.09 | 705.57 | 69.00 |
| LQ | 4.5 | 0.85 | 0.78 | 0.08 | 0.16 | 49449.92 | 1311.39 | 20.00 |
| LQ | 1 | 0.84 | 0.79 | 0.08 | 0.15 | 49472.42 | 1333.90 | 21.00 |
| LQ | 1.5 | 0.84 | 0.79 | 0.08 | 0.15 | 49472.42 | 1333.90 | 21.00 |
| LQ | 2 | 0.84 | 0.79 | 0.08 | 0.15 | 49472.42 | 1333.90 | 21.00 |
| LQ | 2.5 | 0.84 | 0.79 | 0.08 | 0.15 | 49472.42 | 1333.90 | 21.00 |
| LQ | 3 | 0.84 | 0.79 | 0.08 | 0.15 | 49472.42 | 1333.90 | 21.00 |
| LQ | 3.5 | 0.84 | 0.78 | 0.08 | 0.16 | 49472.42 | 1333.90 | 21.00 |
| LQ | 4 | 0.84 | 0.78 | 0.08 | 0.16 | 49472.42 | 1333.90 | 21.00 |
| LQ | 5 | 0.84 | 0.79 | 0.08 | 0.15 | 49512.36 | 1373.84 | 19.00 |
| L | 5 | 0.82 | 0.79 | 0.05 | 0.14 | 50331.13 | 2192.61 | 15.00 |
| L | 1 | 0.82 | 0.79 | 0.05 | 0.14 | 50405.71 | 2267.19 | 15.00 |
| L | 1.5 | 0.82 | 0.79 | 0.05 | 0.14 | 50405.71 | 2267.19 | 15.00 |
| L | 2 | 0.82 | 0.79 | 0.05 | 0.14 | 50405.71 | 2267.19 | 15.00 |
| L | 2.5 | 0.82 | 0.79 | 0.05 | 0.14 | 50405.71 | 2267.19 | 15.00 |
| L | 3 | 0.82 | 0.79 | 0.05 | 0.14 | 50405.71 | 2267.19 | 15.00 |
| L | 3.5 | 0.82 | 0.79 | 0.05 | 0.14 | 50405.71 | 2267.19 | 15.00 |
| L | 4 | 0.82 | 0.79 | 0.05 | 0.14 | 50405.71 | 2267.19 | 15.00 |
| L | 4.5 | 0.81 | 0.79 | 0.05 | 0.14 | 50408.30 | 2269.78 | 15.00 |

L = linear; Q = quadratic; H = hinge; RM = regularization multiplier; AUC = area under the curve; OR = omission rate.

**Supplemental table for Figure 2.**
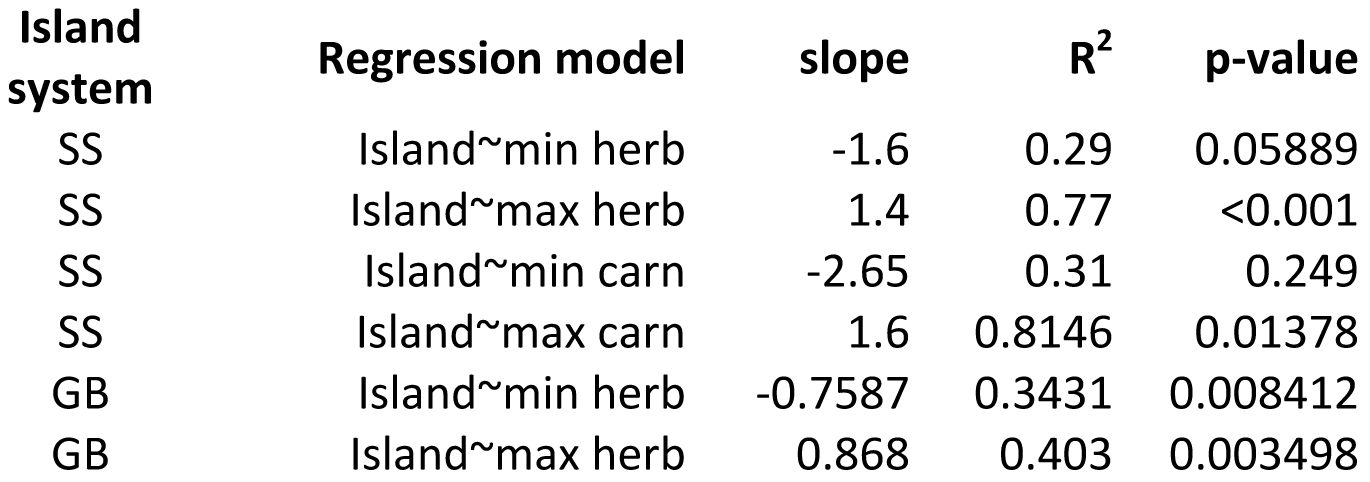
Statistics for the regressions between island area and species’ maximum and minimum body sizes, by trophic levels for the Great Basin and Sunda Shelf mammal fauna. Data for the Great Basin sky islands are from (Brown 1978; Grayson and Livingston 1993) and Sunda Shelf (Okie and Brown 2009).

## References

Åberg, J., Jansson, G., Swenson, J.E. & Angelstam, P. (1995). The effect of matrix on the occurrence of hazel grouse (Bonasa bonasia) in isolated habitat fragments. Oecologia, 103, 265–269

Agrawal, A.A., Ackerly, D.D., Adler, F., Arnold, A.E., Cáceres, C., Doak, D.F., et al. (2007). Filling key gaps in population and community ecology. Frontiers in Ecology and the Environment, 5, 145–152

Altermatt, F., Bieger, A., Carrara, F., Rinaldo, A. & Holyoak, M. (2011). Effects of Connectivity and Recurrent Local Disturbances on Community Structure and Population Density in Experimental Metacommunities. PLOS ONE, 6, e19525

Anderson, C.D., Epperson, B.K., Fortin, M.-J., Holderegger, R., James, P., Rosenberg, M.S., et al. (2010). Considering spatial and temporal scale in landscape-genetic studies of gene flow. Molecular Ecology, 19, 3565–3575

Anderson, R.P. (2013). A framework for using niche models to estimate impacts of climate change on species distributions. Ann. N.Y. Acad. Sci., 1297, 8–28

Anderson, R.P. (2017). When and how should biotic interactions be considered in models of species niches and distributions? Journal of Biogeography, 44, 8–17

Baum, K.A., Haynes, K.J., Dillemuth, F.P. & Cronin, J.T. (2004). The matrix enhances the effectiveness of corridors and stepping stones. Ecology, 85, 2671–2676

Bennett, K.D. (1990). Milankovitch cycles and their effects on species in ecological and evolutionary time. Paleobiology, 16, 11–21

Berger, A. (1988). Milankovitch theory and climate. Reviews of geophysics, 26, 624–657

Bierregaard, R.O., Lovejoy, T.E., Kapos, V., Santos, A.A. dos & Hutchings, R.W. (1992). The Biological Dynamics of Tropical Rainforest Fragments. BioScience, 42, 859–866

Blueweiss, L., Fox, H., Kudzma, V., Nakashima, D., Peters, R. & Sams, S. (1978). Relationships between body size and some life history parameters. Oecologia, 37, 257–272

Bolger, D.T., Alberts, A.C., Sauvajot, R.M., Potenza, P., McCalvin, C., Tran, D., et al. (1997). Response of rodents to habitat fragmentation in coastal southern California. Ecological Applications, 7, 552–563

Bolker, B., Holyoak, M., Křivan, V., Rowe, L. & Schmitz, O. (2003). Connecting theoretical and empirical studies of trait-mediated interactions. Ecology, 84, 1101–1114

Bowman, J., Jaeger, J.A. & Fahrig, L. (2002). Dispersal distance of mammals is proportional to home range size. Ecology, 83, 2049–2055

Boyer, A.G. & Jetz, W. (2014). Extinctions and the loss of ecological function in island bird communities. Global Ecology and Biogeography, 23, 679–688

Brown, J.H. (1971). Mammals on Mountaintops: Nonequilibrium Insular Biogeography. The American Naturalist, 105, 467–478

Brown, J.H., Gillooly, J.F., Allen, A.P., Savage, V.M. & West, G.B. (2004). Toward a Metabolic Theory of Ecology. Ecology, 85, 1771–1789

Brown, J.H. & Kodric-Brown, A. (1977). Turnover rates in insular biogeography: effect of immigration on extinction. Ecology, 58, 445–449

Brown, J.L. (2014). SDMtoolbox: a python-based GIS toolkit for landscape genetic, biogeographic and species distribution model analyses. Methods in Ecology and Evolution, 5, 694–700

Bruno, J.F., Stachowicz, J.J. & Bertness, M.D. (2003). Inclusion of facilitation into ecological theory. Trends in Ecology & Evolution, 18, 119–125

Burger, J.R., Weinberger, V.P. & Marquet, P.A. (2017). Extra-metabolic energy use and the rise in human hyper-density. Scientific Reports, 7

Butaye, J., Jacquemyn, H. & Hermy, M. (2001). Differential colonization causing non-random forest plant community structure in a fragmented agricultural landscape. Ecography, 24, 369–380

Calder, W.A. (1984). Size, function, and life history. Courier Corporation

Carbone, C. & Gittleman, J.L. (2002). A Common Rule for the Scaling of Carnivore Density. Science, 295, 2273–2276

Carlquist, S. (1966). The biota of long-distance dispersal. I. Principles of dispersal and evolution. The Quarterly Review of Biology, 41, 247–270

Chase, J.M. (2003). Community assembly: when should history matter? Oecologia, 136, 489–498

Cody, M.L. & Diamond, J.M. (1975). Ecology and evolution of communities. Harvard University Press

Collinge, S.K. (2000). Effects of Grassland Fragmentation on Insect Species Loss, Colonization, and Movement Patterns. Ecology, 81, 2211–2226

Constable, H., Guralnick, R., Wieczorek, J., Spencer, C., Peterson, A.T. & Committee, TVS. (2010). VertNet: A New Model for Biodiversity Data Sharing. PLOS Biology, 8, e1000309

Crespi, E.J., Rissler, L.J. & Browne, R.A. (2003). Testing Pleistocene refugia theory: phylogeographical analysis of Desmognathus wrighti, a high-elevation salamander in the southern Appalachians. Molecular Ecology, 12, 969–984

Cristofoli, S., Piqueray, J., Dufrêne, M., Bizoux, J.-P. & Mahy, G. (2010). Colonization Credit in Restored Wet Heathlands. Restoration Ecology, 18, 645–655

Crooks, K.R. (2002). Relative sensitivities of mammalian carnivores to habitat fragmentation. Conservation Biology, 16, 488–502

Crooks K.R., et al. (2017) Quantification of habitat fragmentation reveals extinction risk in terrestrial mammals. Proceedings of the National Academy of Sciences. 114: 7635–7640.

Damuth, J. (1981). Population density and body size in mammals. Nature, 290, 699–700

Damuth, J. (1987). Interspecific allometry of population density in mammals and other animals: the independence of body mass and population energy-use. Biological Journal of the Linnean Society, 31, 193–246

Dawson, T.P., Jackson, S.T., House, J.I., Prentice, I.C. & Mace, G.M. (2011). Beyond predictions: biodiversity conservation in a changing climate. science, 332, 53–58

Diffendorfer, J.E., Gaines, M.S. & Holt, R.D. (1995). Habitat fragmentation and movements of three small mammals (Sigmodon, Microtus, and Peromyscus). Ecology, 76, 827–839

Dirnböck, T., Essl, F. & Rabitsch, W. (2011). Disproportional risk for habitat loss of high-altitude endemic species under climate change. Global Change Biology, 17, 990–996

Engemann, K., Sandel, B., Boyle, B., Enquist, B.J., Jørgensen, P.M., Kattge, J., et al. (2016). A plant growth form dataset for the New World. Ecology, 97, 3243–3243

Engler, R., Randin, C.F., Vittoz, P., Czáka, T., Beniston, M., Zimmermann, N.E., et al. (2009). Predicting future distributions of mountain plants under climate change: does dispersal capacity matter? Ecography, 32, 34–45

Enquist, B.J., Norberg, J., Bonser, S.P., Violle, C., Webb, C.T., Henderson, A., et al. (2015). Chapter Nine - Scaling from Traits to Ecosystems: Developing a General Trait Driver Theory via Integrating Trait-Based and Metabolic Scaling Theories. In: Advances in Ecological Research, Trait-Based Ecology - From Structure to Function (ed. Samraat Pawar, G.W. and A.I.D.). Academic Press, pp. 249–318

Estes, J.A., Terborgh, J., Brashares, J.S., Power, M.E., Berger, J., Bond, W.J., et al. (2011). Trophic downgrading of planet Earth. science, 333, 301–306

Estrada, A., Meireles, C., Morales-Castilla, I., Poschlod, P., Vieites, D., Araújo, M.B., et al. (2015). Species’ intrinsic traits inform their range limitations and vulnerability under environmental change. Global Ecology and Biogeography, 24, 849–858

Fahrig, L. (2003). Effects of habitat fragmentation on biodiversity. Annual review of ecology, evolution, and systematics, 34, 487–515

Floyd, C.H., Van Vuren, D.H. & May, B. (2005). Marmots on Great Basin mountaintops: using genetics to test a biogeographic paradigm. Ecology, 86, 2145–2153

Fristoe, T.S. (2015). Energy use by migrants and residents in North American breeding bird communities. Global Ecology and Biogeography, 24, 406–415

Fukami, T. (2015). Historical contingency in community assembly: integrating niches, species pools, and priority effects. Annual Review of Ecology, Evolution, and Systematics, 46, 1–23

Fukami, T. & Nakajima, M. (2011). Community assembly: alternative stable states or alternative transient states? Ecology letters, 14, 973–984

Grayson, D.K. & Livingston, S.D. (1993). Missing Mammals on Great Basin Mountains: Holocene Extinctions and Inadequate Knowledge. Conservation Biology, 7, 527–532

Haddad, N.M., Brudvig, L.A., Clobert, J., Davies, K.F., Gonzalez, A., Holt, R.D., et al. (2015). Habitat fragmentation and its lasting impact on Earth’s ecosystems. Science Advances, 1, e1500052

Hamilton, T.H. (1961). The adaptive significances of intraspecific trends of variation in wing length and body size among bird species. Evolution, 15, 180–195

Hampe, A. & Petit, R.J. (2005). Conserving biodiversity under climate change: the rear edge matters. Ecology letters, 8, 461–467

Hanski, I. (1999). Metapopulation ecology. Oxford University Press

Hanski, I. & Ovaskainen, O. (2003). Metapopulation theory for fragmented landscapes. Theoretical population biology, 64, 119–127

Harrison, S. & Cornell, H. (2008). Toward a better understanding of the regional causes of local community richness. Ecology letters, 11, 969–979

Heaney, L.R. (2000). Dynamic disequilibrium: a long-term, large-scale perspective on the equilibrium model of island biogeography. Global Ecology and Biogeography, 9, 59–74

Heaney, L.R. (2007). Is a new paradigm emerging for oceanic island biogeography? Journal of Biogeography, 34, 753–757

Hennemann, W.W. (1983). Relationship among body mass, metabolic rate and the intrinsic rate of natural increase in mammals. Oecologia, 56, 104–108

Hijmans, R.J., Cameron, S.E., Parra, J.L., Jones, P.G. & Jarvis, A. (2005). Very high resolution interpolated climate surfaces for global land areas. Int. J. Climatol., 25, 1965–1978

Hijmans, R.J., Phillips, S., Leathwick, J. & Elith, J. (2015). dismo: Species distribution modeling. R package version 1.0-12. The R Foundation for Statistical Computing, Vienna http://cran.rproject.org

Holt, R.D. (2009). Toward a Trophic Island Biology. In: The Theory of Island Biogeography Revisited. Princeton University Press, pp. 143–185

Holt, R.D., Robinson, G.R. & Gaines, M.S. (1995). Vegetation dynamics in an experimentally fragmented landscape. Ecology, 76, 1610–1624

Holyoak, M. (2014). Connectance and Connectivity. In: Encyclopedia of Ecology. Newnes

Hunter, M.D. (2002). Landscape structure, habitat fragmentation, and the ecology of insects. Agricultural and Forest Entomology, 4, 159–166

Jackson, S.T., Betancourt, J.L., Booth, R.K. & Gray, S.T. (2009). Ecology and the ratchet of events: climate variability, niche dimensions, and species distributions. Proceedings of the National Academy of Sciences, 106, 19685–19692

Jackson, S.T. & Sax, D.F. (2010). Balancing biodiversity in a changing environment: extinction debt, immigration credit and species turnover. Trends in ecology & evolution, 25, 153–160

Jacquet, C., Mouillot, D., Kulbicki, M. & Gravel, D. (2017). Extensions of Island Biogeography Theory predict the scaling of functional trait composition with habitat area and isolation. Ecol Lett, 20, 135–146

Jones, K.E., Bielby, J., Cardillo, M., Fritz, S.A., O’Dell, J., Orme, C.D.L., et al. (2009). PanTHERIA: species-level database of life history, ecology, and geography of extant and recently extinct mammals. Ecology, 90, 2648–2648

Kearney, M. & Porter, W. (2009). Mechanistic niche modelling: combining physiological and spatial data to predict species’ ranges. Ecology Letters, 12, 334–350

Kelt, D.A. & Van Vuren, D.H. (2001). The ecology and macroecology of mammalian home range area. The American Naturalist, 157, 637–645

Keymer, J.E., Marquet, P.A., Velasco-Hernández, J.X. & Levin, S.A. (2000). Extinction Thresholds and Metapopulation Persistence in Dynamic Landscapes. The American Naturalist, 156, 478–494

Kidwell, S.M. (2013). Time-averaging and fidelity of modern death assemblages: building a taphonomic foundation for conservation palaeobiology. Palaeontology, 56, 487–522

Kirmer, A., Tischew, S., Ozinga, W.A., Von Lampe, M., Baasch, A. & Van Groenendael, J.M. (2008). Importance of regional species pools and functional traits in colonization processes: predicting re-colonization after large-scale destruction of ecosystems. Journal of Applied Ecology, 45, 1523–1530

Kitzes, J. & Harte, J. (2015). Predicting extinction debt from community patterns. Ecology, 96, 2127–2136

Larsen, T.H., Williams, N.M. & Kremen, C. (2005). Extinction order and altered community structure rapidly disrupt ecosystem functioning. Ecology letters, 8, 538–547

Lasky, J.R., Sun, I., Su, S.-H., Chen, Z.-S. & Keitt, T.H. (2013). Trait-mediated effects of environmental filtering on tree community dynamics. Journal of Ecology, 101, 722–733

Laurance, W.F. (2008). Theory meets reality: how habitat fragmentation research has transcended island biogeographic theory. Biological conservation, 141, 1731–1744

Laurance, W.F., Lovejoy, T.E., Vasconcelos, H.L., Bruna, E.M., Didham, R.K., Stouffer, P.C., et al. (2002). Ecosystem Decay of Amazonian Forest Fragments: a 22-Year Investigation. Conservation Biology, 16, 605–618

Leibold, M.A., Holyoak, M., Mouquet, N., Amarasekare, P., Chase, J.M., Hoopes, M.F., et al. (2004). The metacommunity concept: a framework for multi-scale community ecology. Ecology Letters, 7, 601–613

Leimu, R. & Fischer, M. (2008). A meta-analysis of local adaptation in plants. PloS one, 3, e4010

Lenzner, B., Weigelt, P., Kreft, H., Beierkuhnlein, C. & Steinbauer, M.J. (2017). The general dynamic model of island biogeography revisited at the level of major flowering plant families. Journal of Biogeography, 44, 1029–1040

Lessa, E.P. & Farina, R.A. (1996). Reassessment of extinction patterns among the late Pleistocene mammals of South America. Palaeontology, 39, 651–662

Lessard, J.-P., Weinstein, B.G., Borregaard, M.K., Marske, K.A., Martin, D.R., McGuire, J.A., et al. (2016). Process-based species pools reveal the hidden signature of biotic interactions amid the influence of temperature filtering. The American Naturalist, 187, 75–88

Lomolino, M.V. (1993). Winter filtering, immigrant selection and species composition of insular mammals of Lake Huron. Ecography, 16, 24–30

MacArthur, R.H. & Wilson, E.O. (1963). An equilibrium theory of insular zoogeography. Evolution, 373–387

MacArthur, R.H. & Wilson, E.O. (1967). The theory of island biogeography. Monographs in population biology

Manning, A.D., Fischer, J., Felton, A., Newell, B., Steffen, W. & Lindenmayer, D.B. (2009). Landscape fluidity–a unifying perspective for understanding and adapting to global change. Journal of Biogeography, 36, 193–199

Marquet, P.A. (2002). Of predators, prey, and power laws. Science, 295, 2229–2230

Marquet, P.A., Allen, A.P., Brown, J.H., Dunne, J.A., Enquist, B.J., Gillooly, J.F., et al. (2014). On theory in ecology. BioScience, 64, 701–710

Marquet, P.A. & Taper, M.L. (1998). On size and area: Patterns of mammalian body size extremes across landmasses. Evolutionary Ecology, 12, 127–139

May, R.M., Levin, S.A. & Sugihara, G. (2008). Complex systems: Ecology for bankers. Nature, 451, 893–895

McDonald, K.A. & Brown, J.H. (1992). Using montane mammals to model extinctions due to global change. Conservation Biology, 6, 409–415

McGill, B.J., Enquist, B.J., Weiher, E. & Westoby, M. (2006). Rebuilding community ecology from functional traits. Trends in Ecology & Evolution, 21, 178–185

McIntire, E.J., Schultz, C.B. & Crone, E.E. (2007). Designing a network for butterfly habitat restoration: where individuals, populations and landscapes interact. Journal of Applied Ecology, 44, 725–736

McRae, B.H., Dickson, B.G., Keitt, T.H. & Shah, V.B. (2008). Using circuit theory to model connectivity in ecology, evolution, and conservation. Ecology, 89, 2712–2724

Meijaard, E. (2001). Successful sea-crossing by land mammals; a matter of luck, and a big body: a preliminary and simplified model. Geological Research and Development Centre’s Special Publication, 27, 87–92

Milankovitch, M. (1941). History of radiation on the Earth and its use for the problem of the ice ages. K. Serb. Akad. Beogr

Moore, R.P., Robinson, W.D., Lovette, I.J. & Robinson, T.R. (2008). Experimental evidence for extreme dispersal limitation in tropical forest birds. Ecology letters, 11, 960–968

Okie, J.G. & Brown, J.H. (2009). Niches, body sizes, and the disassembly of mammal communities on the Sunda Shelf islands. Proceedings of the National Academy of Sciences, 106, 19679–19684

Oksanen, L., Fretwell, S.D., Arruda, J. & Niemela, P. (1981). Exploitation Ecosystems in Gradients of Primary Productivity. The American Naturalist, 118, 240–261

Olszewski, T. (1999). Taking advantage of time-averaging. Paleobiology, 25, 226–238

Patino, J., Whittaker, R.J., Borges, P.A., Fernández-Palacios, J.M., Ah-Peng, C., Araújo, M.B., et al. (2017). A roadmap for island biology: 50 fundamental questions after 50 years of The Theory of Island Biogeography. Journal of Biogeography, 44, 963–983

Patterson, B.D. (1987). The Principle of Nested Subsets and Its Implications for Biological Conservation. Conservation Biology, 1, 323–334

Peters, R.H. (1986). The ecological implications of body size. Cambridge University Press

Peterson, A.T. (2011). Ecological Niches and Geographic Distributions (MPB-49). Princeton University Press

Peterson, A.T. & Ammann, C.M. (2013). Global patterns of connectivity and isolation of populations of forest bird species in the late Pleistocene. Global Ecology and Biogeography, 22, 596–606

Prates, I., Xue, A.T., Brown, J.L., Alvarado-Serrano, D.F., Rodrigues, M.T., Hickerson, M.J., et al. (2016). Inferring responses to climate dynamics from historical demography in neotropical forest lizards. Proceedings of the National Academy of Sciences, 113, 7978–7985

Prevedello, J.A. & Vieira, M.V. (2010). Does the type of matrix matter? A quantitative review of the evidence. Biodiversity and Conservation, 19, 1205–1223

Pulliam, H.R. (1988). Sources, sinks, and population regulation. The American Naturalist, 132, 652–661

Rahmstorf, S. (2007). A semi-empirical approach to projecting future sea-level rise. Science, 315, 368–370

Rickart, E.A. (2001). Elevational diversity gradients, biogeography and the structure of montane mammal communities in the intermountain region of North America. Global Ecology and Biogeography, 10, 77–100

Ricketts, T.H. (2001). The matrix matters: effective isolation in fragmented landscapes. The American Naturalist, 158, 87–99

Rowe, R.J. (2005). Elevational gradient analyses and the use of historical museum specimens: a cautionary tale. Journal of Biogeography, 32, 1883–1897

Rowe, R.J. (2009). Environmental and geometric drivers of small mammal diversity along elevational gradients in Utah. Ecography, 32, 411–422

Roy, K., Valentine, J.W., Jablonski, D. & Kidwell, S.M. (1996). Scales of climatic variability and time averaging in Pleistocene biotas: implications for ecology and evolution. Trends in Ecology & Evolution, 11, 458–463

Savage, V.M., Gillooly, J.F., Brown, J.H., West, G.B. & Charnov, E.L. (2004). Effects of body size and temperature on population growth. The American Naturalist, 163, 429–441

Scheffer, M., Carpenter, S., Foley, J.A., Folke, C. & Walker, B. (2001). Catastrophic shifts in ecosystems. Nature, 413, 591–596

Seidler, T.G. & Plotkin, J.B. (2006). Seed Dispersal and Spatial Pattern in Tropical Trees. PLOS Biology, 4, e344

Shaffer, M.L. (1981). Minimum population sizes for species conservation. BioScience, 31, 131–134

Sibly, R.M., Brown, J.H. & Kodric-Brown, A. (2012). Metabolic Ecology: A Scaling Approach. John Wiley & Sons

Silva, M. & Downing, J.A. (1995). The Allometric Scaling of Density and Body Mass: A Nonlinear Relationship for Terrestrial Mammals. The American Naturalist, 145, 704–727

Simberloff, D.S. & Wilson, E.O. (1969). Experimental zoogeography of islands: the colonization of empty islands. Ecology, 50, 278–296

Soley-Guardia, M., Gutiérrez, E.E., Thomas, D.M., Ochoa-G, J., Aguilera, M. & Anderson, R.P. (2016). Are we overestimating the niche? Removing marginal localities helps ecological niche models detect environmental barriers. Ecology and evolution

Soulé, M.E. (1987). Viable populations for conservation. Cambridge university press

Soulé, M.E., Alberts, A.C. & Bolger, D.T. (1992). The effects of habitat fragmentation on chaparral plants and vertebrates. Oikos, 39–47

Steinbauer, M.J., Dolos, K., Field, R., Reineking, B. & Beierkuhnlein, C. (2013). Re-evaluating the general dynamic theory of oceanic island biogeography. frontiers of biogeography, 5, 185–194

Steinbauer, M.J., Otto, R., Naranjo-Cigala, A., Beierkuhnlein, C. & Fernández-Palacios, J.-M. (2012). Increase of island endemism with altitude–speciation processes on oceanic islands. Ecography, 35, 23–32

Stephens, P.A. (2016). Population viability analysis. Oxford University Press

Stouffer, P.C. & Bierregaard, R.O. (1995). Use of Amazonian forest fragments by understory insectivorous birds. Ecology, 76, 2429–2445

Tamburello, N., Côté, I.M. & Dulvy, N.K. (2015). Energy and the scaling of animal space use. The American Naturalist, 186, 196–211

Terborgh, J., Estes, J.A., Paquet, P., Ralls, K., Boyd-Herger, D., Miller, B.J., et al. (1999). The role of top carnivores in regulating terrestrial ecosystems

Tilman, D., May, R.M., Lehman, C.L. & Nowak, M.A. (1994). Habitat destruction and the extinction debt

Urban, D. & Keitt, T. (2001). Landscape Connectivity: A Graph-Theoretic Perspective. Ecology, 82, 1205–1218

Vannette, R.L. & Fukami, T. (2014). Historical contingency in species interactions: towards niche-based predictions. Ecology Letters, 17, 115–124

Violle, C., Reich, P.B., Pacala, S.W., Enquist, B.J. & Kattge, J. (2014). The emergence and promise of functional biogeography. PNAS, 111, 13690–13696

Waltari, E. & Guralnick, R.P. (2009). Ecological niche modelling of montane mammals in the Great Basin, North America: examining past and present connectivity of species across basins and ranges. Journal of Biogeography, 36, 148–161

Warren, B.H., Simberloff, D., Ricklefs, R.E., Aguilée, R., Condamine, F.L., Gravel, D., et al. (2015). Islands as model systems in ecology and evolution: prospects fifty years after MacArthur-Wilson. Ecology Letters, 18, 200–217

Weigelt, P., Steinbauer, M.J., Cabral, J.S. & Kreft, H. (2016). Late Quaternary climate change shapes island biodiversity. Nature, 532, 99–102

Werner, E.E. & Peacor, S.D. (2003). A review of trait-mediated indirect interactions in ecological communities. Ecology, 84, 1083–1100

Whittaker, R.J., Triantis, K.A. & Ladle, R.J. (2008). A general dynamic theory of oceanic island biogeography. Journal of Biogeography, 35, 977–994

Whittaker, R. J., Fernández-Palacios, J. M., Matthews, T. J., Borregaard, M. K. & Triantis, K. A. (2017) Island biogeography: Taking the long view of nature’s laboratories. Science 357

Wiens, J.J., Ackerly, D.D., Allen, A.P., Anacker, B.L., Buckley, L.B., Cornell, H.V., et al. (2010). Niche conservatism as an emerging principle in ecology and conservation biology. Ecology letters, 13, 1310–1324

Wright, D.H. & Reeves, J.H. (1992). On the meaning and measurement of nestedness of species assemblages. Oecologia, 92, 416–428

